# Defining and Controlling Axial Nephron Patterning in Human Kidney Organoids with Synthetic Wnt-Secreting Organizers

**DOI:** 10.1101/2024.11.30.626171

**Authors:** Connor C. Fausto, Fokion Glykofrydis, Navneet Kumar, Jack Schnell, Reka L. Csipan, Faith De Kuyper, Brendan Grubbs, Matthew Thornton, MaryAnne Achieng, Leonardo Morsut, Nils O. Lindström

## Abstract

Current human pluripotent stem cell-derived kidney organoids contain nephron-like structures that lack organotypic patterning. It is thought that during human development, nephrons form their proximal-distal axial polarity in response to collecting duct-derived signals that are absent in kidney organoids. To delineate how nephron polarities establish, we profiled human kidney development by spatial transcriptomic approaches. Our analyses describe a new axial polarity in the nephron and demonstrate that the nephron proximal-distal polarity develops adjacent to a transcriptional boundary in the collecting duct where non-canonical *WNT11* is downregulated and canonical *WNT9B* ligand is upregulated. The nephron region closest to this boundary in turn activates a series of canonical WNT target genes inferring positional nephron identities. To establish whether a canonical WNT source can improve organoid patterning to an *in vivo*-like state, we bioengineered self-organizing WNT-secreting synthetic organizers. Organizer-coupled kidney organoids respond to WNT ligands by forming expression gradients and developing distal cell identities. Tuning the WNT dose produced nephrons with continuous patterning along the proximal-distal axis. Strikingly, polarized iPSC-derived nephrons directed their distal tubules towards the WNT-source, indicating axial patterning and morphogenetic programs are tuned by WNTs from the synthetic organizers. Our data present a strategy to control organ patterning, build an artificial kidney, and highlights the power of synthetic organizer systems for advancing organoid models.

## Introduction

Kidney organoids generated from human induced pluripotent stem cells (hiPSCs) represent a promising source of synthetic kidney tissue to solve the scarcity of donor kidneys available for transplantation. To reach clinical relevance, kidney organoids must generate patterned nephrons that filter blood in renal corpuscles, reabsorb electrolytes in proximal convoluted tubules, regulate water and concentrate urine in the loop-of-Henle and distal tubules prior to draining into a collecting duct. However, current kidney organoids contain nephrons that display limited maturation of distal cell types, and incomplete proximal-distal (PD) patterning. This suggests that organoid derivation methods can be modified to include key mechanisms and tissues responsible for nephron patterning and kidney development, steps that would broaden the potential of organoid nephrons as therapeutics.

Nephron patterning arises during kidney organogenesis ^1–4^. Kidneys develop from reciprocal interactions between the collecting duct (CD) and the metanephric mesenchyme that contains nephron progenitor cells (NPCs) and stromal populations ^3^. Throughout kidney development the forming nephron and CD develop in contiguity (see Fig. 1a for a schematic). The developing CD consists of molecularly distinct tip (*WNT11*^+^) and stalk (*WNT9B*^+^) populations, containing CD progenitors and their progeny, respectively. NPCs marked by SIX2, CITED1, MEOX1, and PCDH15, are found surrounding CD tips and undergo balanced self-renewal and differentiation in response to CD stalk-derived WNT9B ligands that activate β-catenin signaling in NPCs to initiate nephrogenesis ^5–12^. Nephron formation is first observed as a stream of NPCs that exit their tip niche, downregulate *SIX2* and condensate into pre-tubular aggregates (PTA; expressing *WNT4*, *FGF8* and *LHX1*) under each CD tip ^13–15^. The PTA develops into a lumenized renal vesicle (RV) with polarized *JAG1* and *CDH1* expression towards the CD stalk as more NPCs are gradually recruited to the opposite end of the RV ^15^. The RV thereafter develops into Comma- and S-shaped body nephrons (CSB and SSBs, respectively) that adopt distal (*POU3F3*, *TFAP2A*, *MECOM*, *SOX9*, *CDH1*), medial (*IRX1*, *JAG1*), proximal (*HNF4A*) and podocyte (*MAFB*, *WT1*) segments; these identities reflect their relative time of arrival into the forming nephron ^15–19^. Our studies in mice have shown that the distal nephron is marked by high β-catenin activity and manipulation of Wnt/β-catenin signaling leads to expansion of distal segments at the expense of medial and proximal segments ^20^. Given the spatial relationship between the forming nephron and the CD, and Wnt and β-catenin components controlling distal nephron development and nephrogenesis, it has been hypothesized that the Wnt-expressing CD serves as a signaling organizer for the nephron. Whether this holds true in the developing human kidney remains unknown.

**Figure 1.**
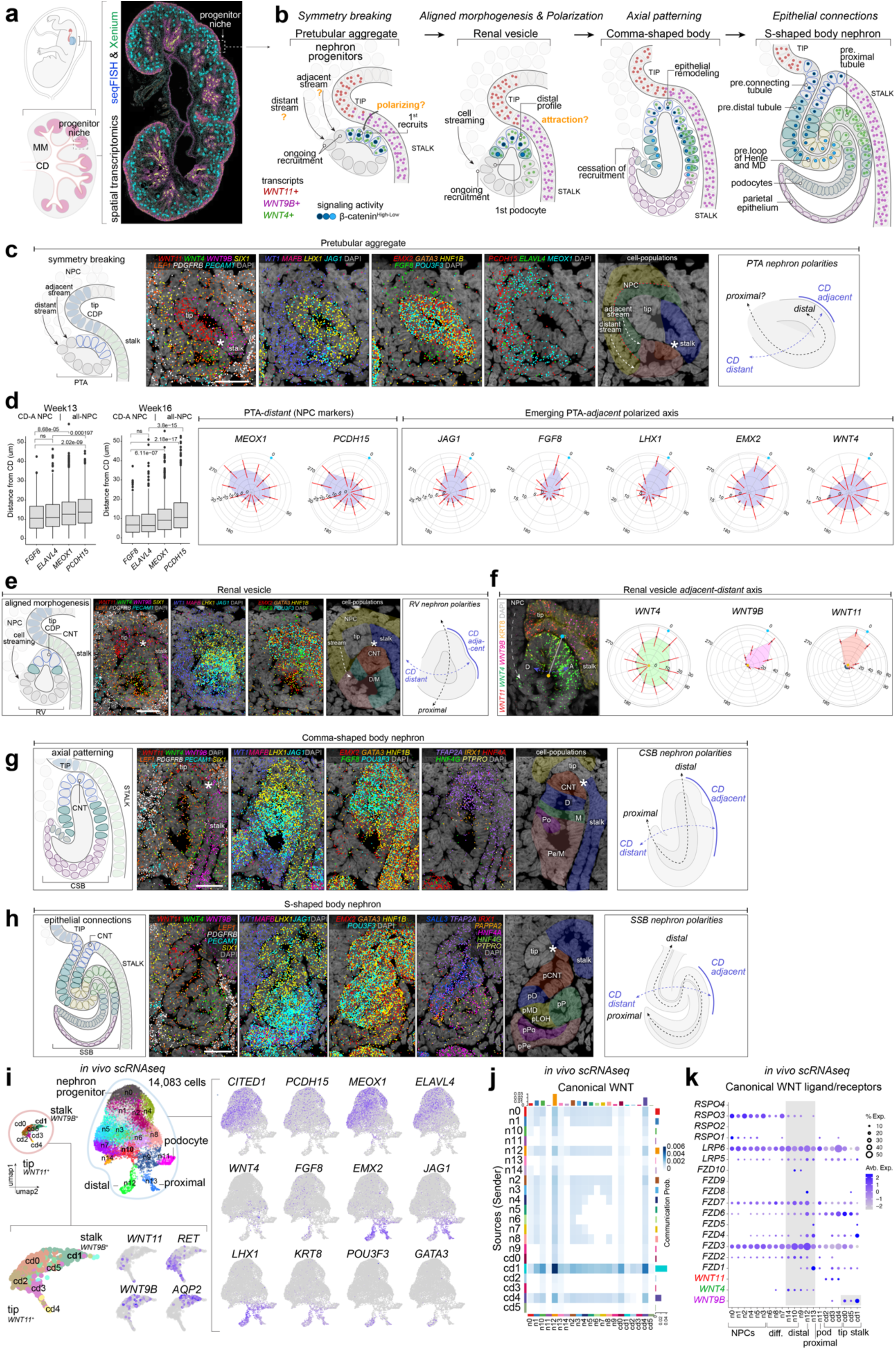
Human nephrons develop multi-axial patterning and morphogenesis relative to collecting duct-localized WNT ligands. **a,** Spatial transcriptomic analyses of developing human kidneys and nephrogenesis (MM: metanephric mesenchyme, CD: collecting duct). **b,** Schematic of nephrogenesis and CD expressed WNT ligands (*WNT11*: CD tip; *WNT9B*: CD stalk) and nephron expressed *WNT4*, proposed β-catenin signaling, and cell populations based on coloring and labels. **c,** Transcript distribution at the pretubular aggregate (PTA) stage (middle panels), schematics showing anatomies (left), inferred cell-populations and CD adjacent-distant (AD) and proximal-distal (PD) axes (right). **d,** Graphs showing distance between the CD surface and transcripts in nephron progenitors (NPCs) at week 13 and 16 (left; CD adjacent: CD-A; CD-distant: CD-D) and transcript distribution across CD-A to CD-D axis in the PTA (right). **e,** Transcript distribution at the renal vesicle (RV) stage with schematics as in (c). **f,** *WNT4* transcript polarization in RVs relating to *WNT11* and *WNT9B* with radar plots depicting transcript frequency distribution (cyan dot: *WNT11/WNT9B* transition; yellow dot: center of RV; red lines: standard deviation). **g-h,** Transcript distribution at the comma-shaped (CSB) and S-shaped body (SSB) stages with schematics outlining cell populations, and PD and AD axes. Scale bars in c-h are 25 µm. **i,** UMAP plot of scRNAseq data containing CD clusters (cd0-5), and nephrogenic NPC to RV clusters (n0-14) with cell markers shown as feature plots matching UMAP annotation. **j.** Plot showing probability of canonical WNT receptor (x-axis)-ligand (y-axis) pairing. **k,** Dotplot showing expression of WNT ligands, Frizzled receptors, LRP co-receptors, and R-spondin potentiators in early nephrogenesis.

Several kidney organoid models describe how to generate kidney cell-types by mimicking a developmental progression from pluripotency to metanephric mesenchyme-like cell states ^21–24^. Their strengths are in their relatively rapid timeline. Nephrons form between 10 and 15 days from cells leaving pluripotency. However, off-target cell-populations are frequently reported, and the forming nephrons display limited maturation and function compared to adult *in vivo* nephrons ^18,25,26^. These organoid models typically lack a CD epithelium which can be derived from an anterior intermediate mesoderm lineage through separate differentiation protocols ^27–29^. Instead, nephron differentiation is induced by pulsing the organoids with a WNT/β-catenin agonist ^21^. In the Takasato protocol this occurs 3 days prior to the formation of nephron-like structures, and organoid nephrons therefore organize without a CD and its secretome ^21^.

Determining whether kidney organoid nephrons can be locally controlled by CD-derived signals to generate normally axial PD patterning is therefore an important goal. Controlling patterning in other organoid models has been attempted with microfluidics delivery of growth factors and growth factors from cells ^30,31^. Recently, cell-based signaling centers have been used to pattern embryoid bodies (EBs) ^32^ in mESC and hiPSC derived brain organoids ^33^. These foundational studies show that delivery of growth factors from adjacent cell sources can initiate and locally pattern regions of stem-cell-derived constructs. However, whether these approaches can finely pattern human organoids and mimic cellular-scale morphological and patterning features as seen *in vivo*, is not yet determined.

We hypothesized that recapitulating the CD-to-nephron signaling relationship in kidney organoids can normalize organoid nephron patterning. Here we first resolve the spatial transcriptional landscape of the developing human kidney and show nephrons possess a novel axial polarity. We show that nephrons develop adjacent to the transition point where CD precursors exit their *WNT11^+^* niche and differentiate into CD stalk (*WNT9B^+^*). Nephron cells adjacent to this transition express canonical WNT targets, form a distal nephron identity, and ultimately connect to the CD epithelium to form a luminal connection that enables drainage of urine into the CD. Detailing the nephron transcriptional profile demonstrates that the process of nephrogenesis occurs via an inherently asymmetric mechanism. Nephrons first develop a CD-<u>a</u>djacent to CD-<u>d</u>istant (AD) axial polarity that presages nephron anatomies and formation of its known PD axial polarity. Comparative analyses between kidneys and organoids reveal that organoids lack a canonical WNT signal input but can respond to WNT-secreting synthetic organizer cells. Coupling organoid nephrons with synthetic organizers drives a canonical WNT response, promotes distal nephron differentiation programs, and strikingly generates a morphogenetic response in nephrons prompting them to orient their elongating axes towards the WNT source. Our data highlight a strategy to engineer axial asymmetry and patterning in kidney organoids, control differentiation programs, and pave the way towards artificial kidneys that connect to an artificial epithelium and drain urine.

## Results

### Cross-lineage communication correlates with nephron polarization and morphogenesis in the human developing kidney

To scrutinize signaling interactions within the forming human nephron and its niche, we performed spatial transcriptional analyses using Xenium and seqFISH ^34,35^, mapping the expression of hundreds of genes at two stages of kidney development rich in nephron formation (**Fig.1**, **Fig.S1**). We analyzed the expression of known and putative WNT target genes in relationship to WNT ligands and markers of cell types present in the developing kidney (**Fig.1a**). Given that nephrons develop asynchronously and continuously across the developing kidney cortex, this allowed us to simultaneously observe all stages of early nephrogenesis. Our observations, described below, led us to hypothesize that the WNT expressing interface of the CD functions as a *de facto* organizer of the forming nephron during its aggregation and formation of the nephron axial polarity (**Fig. 1b**).

The first stage of nephrogenesis, the PTA, is an approximately 75 µm wide cell-aggregate which forms within 3 to 4 cell distances of the point where CD progenitors (*WNT11^+^*) transition into CD stalk (*WNT9B*^+^); marked with an asterisk in Fig.1c. We have previously shown that mesenchymal NPCs migrate and gradually integrate into the forming PTA and RV ^15^. Here, we noticed that NPCs outside the PTA demonstrate polarized abundance of transcripts (**Fig.S2a**). A CD-adjacent stream of NPCs (*ELAVL4*^high^/*FGF8*^high^), and a CD-distant stream (*ELAVL4*^low^/*FGF8*^low^) can be distinguished compared to other NPC markers (*PCDH15* and *MEOX1*) that are expressed equally in the CD-adjacent and CD-distant NPCs (**Fig.1d**; **Fig.S2b**). The CD-adjacent stream connects to the PTA immediately next to the CD while the CD-distant stream connects to the PTA one cell layer away from the CD. The distribution bias for *FGF8* and *ELAVL4* is evident in both week 13 and 16 kidneys (**Fig. 1d**). The distribution of *ELAVL4* and *FGF8* become progressively more biased towards the CD-adjacent NPCs between week 13 and 16, while *MEOX1* and *PCDH15* remain equal in CD-adjacent and CD-distant NPCs. The broader distribution is likely a consequence of the thicker NPC layers present at week 13 ^10,11^ that thin during development. The local maxima of *FGF8* and *ELAVL4* near the CD-epithelium indicate that their expression is induced in response to a polarized CD-derived signal (**Fig.1d**, **Fig.S2a,b**). Cells at the site of active recruitment in both streams (indicated as the end of the dashed arrows in Fig.1c, right most spatial transcriptomic panel) are *WNT4^low^*, forming further evidence for the PTA being asymmetric. These observations support a scenario by which the 2 streams of nephron progenitors form conveyor-belts of cells that set up an intrinsic asymmetry in the epithelializing nephron polarized along a ‘*CD adjacent’* to ‘*CD distant’* axis.

The PTA consists of a group of cells morphologically distinct from NPCs. PTAs demonstrate distinctly polarized gene expression depending on cells’ position relative to the *WNT11/WNT9B* transition point. The cells of the PTA closest to the *WNT11/WNT9B* transition point and along the *WNT9B*^+^ CD-adjacent surface of the PTA upregulate WNT-β-catenin targets *FGF8*, *LHX1*, *JAG1*, *EMX2*, *WNT4* whereas NPC marker genes *PCDH15*, and *MEOX1* are gradually reduced in CD-adjacent cells (**Fig.1c-d**). These data indicate the PTA forms asymmetrically and from the outset contains cell subdivisions. PTAs progress to develop into RVs, epithelial pear-shaped cysts with an oblong distal domain extending in the direction of the elbow of the CD stalk, the whole structure approximately 100 µm long. At this stage, the NPC stream appears as a single file of cells that connects with the proximal CD-distant end of the RV, s the distal RV remains in tight association with the CD *WNT11*/*WNT9B* transition point (marked with an asterisk in Fig.1e). While the distal-most region thus remains in a fixed position, gene expression-wise, the RV has grown and patterned, forming a distal region (also known as connecting tubule, *GATA3^+^*), an expanding medial domain (*JAG1^+^, HNF1B^+^*), and a proximal domain (*WT1*^+^), where NPCs are recruited (**Fig.1e**; **Fig.S2c**). The emerging connecting tubule and distal-medial domains are simultaneously patterned along their AD and PD axes, and the CD-adjacent RV surface displays polarized expression of *WNT4, FGF8, LHX1, and EMX2* towards the *WNT9B*^+^ source (**Fig.1e**, **Fig.S2c**).

To better define the CD-adjacent to distant transcriptional asymmetry we spatially mapped *WNT4* expression in the RV relative to the CD tip-stalk transition point. To do this, we performed RNA scope *in situ* hybridization against *WNT4*, *WNT11*, and *WNT9B,* and immunolabeled KRT8 (marks CD and distal nephron) to define cell boundaries and *WNT4* expression. *WNT4* transcripts were asymmetrically distributed towards the *WNT9B^+^/*KRT8*^+^*CD stalk (**Fig.1f**). Quantification of the spatial distribution of *WNT4* puncta shows high *WNT4* near the *WNT9B* source and low *WNT4* in cells further from the *WNT9B* source at the site of NPC recruitment and the CD distant domain (**Fig.1f**). These data confirm the existence of a CD distant-adjacent axis in the RV.

After initial asymmetry forms in the RV, it develops into the CSB (**Fig.1g**; **Fig.S2d**) and thereafter the SSB (**Fig.1h**; **Fig.S2e**) nephron stages. Throughout this process, the nephron remains defined by the *WNT11/WNT9B* transition point in the CD. The CSB nephron elongates along its PD axis and is a transitory stage to the more elaborately patterned SSB. CSB cells nearest the CD *WNT11/WNT9B* transition, now form the connecting tubule precursor domain, which is characterized by downregulation of *WNT4* and *FGF8*, and expression of *GATA3, POU3F3, and TFAP2A*. While some *WNT4* transcripts remain in those cells, the *WNT4* expressing domain shifts proximally along the length of the CD-adjacent nephron abutting the *WNT9B* source. *EMX2*, *TFAP2A*, *POU3F3*, remain high, while genes marking later nephron precursor specification (*IRX1*, *MAFB, WT1*) are emerging or defining small subset cells in the medial, podocyte, and parietal epithelium (**Fig.1g**). Other more mature markers such as PTPRO (podocyte) and HNF4A/G (proximal) have not yet been activated. As the nephron then transitions to the SSB, it adopts new morphological features of convolution, particularly the deep glomerular cleft but also the separation of the distal and medial domains. The SSB is characterized by *WNT4* expression that now only persists in the region most closely adjacent to the WNT9B source. This domain is the *HNF4A/G*^+^ proximal precursor (pP) (**Fig.1h**; **Fig.S2e**). Several putative precursors of specific nephron sub-populations have developed by the SSB stage, marked by expression of (*GATA3*^+^/*SALL3*^+^/*POU3F3*^+^/*TFAP2A*^+^) in the connecting tubule precursors (pCNT) which is now connected to the CD via a patent luminal connection, (*GATA3^l^*^ow^/*TFAP2A*^+^) in distal precursors (pDT), (*IRX1*^+^/*PAPPA2*^+^) in loop of Henle and macula densa precursors (pLOH/pMD), proximal precursors (pP) (*HNF4A/G*^+^/*HNF1B*^+^), parietal epithelium precursors (pPE - not defined in figure), and podocyte precursors (pPod) (*MAFB*^+^/*PTPRO*^+^). These morphogenetic steps are now defined at high resolution via spatial transcriptomic analyses and revealing the stepwise patterning of the developing human nephron.

To provide an independent assessment of the hypothesis that the *WNT11/WNT9B* junction functions as a point of reference and organization for early nephrogenesis, we scrutinized single-cell sequencing data (scRNA-seq) from kidneys temporally matched with our spatial transcriptional data ^16,18^ (**Fig.1i**; **Fig.S3; Supplementary table 1**). We resolved cell-types by iteratively clustering the nephrogenic and collecting duct lineages and identified *WNT11*^+^/*RET*^+^ CD tip progenitors, *WNT9B*^+^/*AQP2*^+^ CD stalk populations, NPCs (*CITED1*^+^/*MEOX1*^+^), PTAs (*WNT4*^+^/*PAX8*^+^), and differentiating RV populations. We queried receptor-ligand interaction via *CellChat* ^36^, which highlights potential pair-wise ligand/receptor interactions between cell populations (**Fig.1j,k**; **Fig.S3d**). The *WNT9B*-expressing CD cell cluster “cd1” is predicted to be the ligand-producing cells that target PTA and RV populations “n10” and “n12”, enriched in expression of FZD receptors, through a canonical WNT signal (**Fig.1j,k**). Consistent with that prediction is the expression of *WNT9B* and *WNT4* in cd1 and n10/12, respectively, setting up the WNT9B driven canonical response (*WNT4*) in the PTA and RV. Receiver cells in n12 correspond to early distal progenitors positive for *POU3F3* and *GATA3*. Based on these analyses, there is a collective view emerging predicting a canonical WNT interaction between *WNT9B*^+^ cells in the CD and *WNT4*^+^ cells in the PTA that is sustained in the RV stage.

Our data therefore suggest that *WNT9B* from the ureteric stalk acts as a short-range morphogen driving gene expression in NPCs, and the PTA and RV. Overall, these spatial transcriptomic analyses of human developing kidneys provide a unique view to the inherent asymmetric transcriptional landscape observed during nephron development and support the hypothesis that *WNT9B*^+^ CD cells function as spatial organizers for the forming nephron with a distal nephron program being initiated in the early nephron adjacent to the WNT9B source.

### Human kidney organoids lacking ureteric epithelium contain non-polarized nephrons

To evaluate nephron patterning in organoids (lacking a CD), we performed time-course immunofluorescence and transcriptomic analyses of organoid nephrogenesis in a kidney hiPSC-derived model adapted from the Takasato protocol ^21,37^ (**Fig.2a**). To generate organoids, hiPSCs are treated with a WNT agonist (GSK3-β inhibitor CHIR99021 (CHIR)) for 5 days, and CHIR and FGF9 until day 7 to sequentially differentiate posterior intermediate mesoderm and metanephric mesenchyme. On day 7, the 2D cell layers are transitioned into 3D discs cultured in transwell filters at the air-liquid interface and pulsed with CHIR for 1 hour, and thereafter maintained in FGF9 to promote cell survival until day 12, after which organoids are kept in basal media to enable nephron formation and patterning.

**Figure 2.**
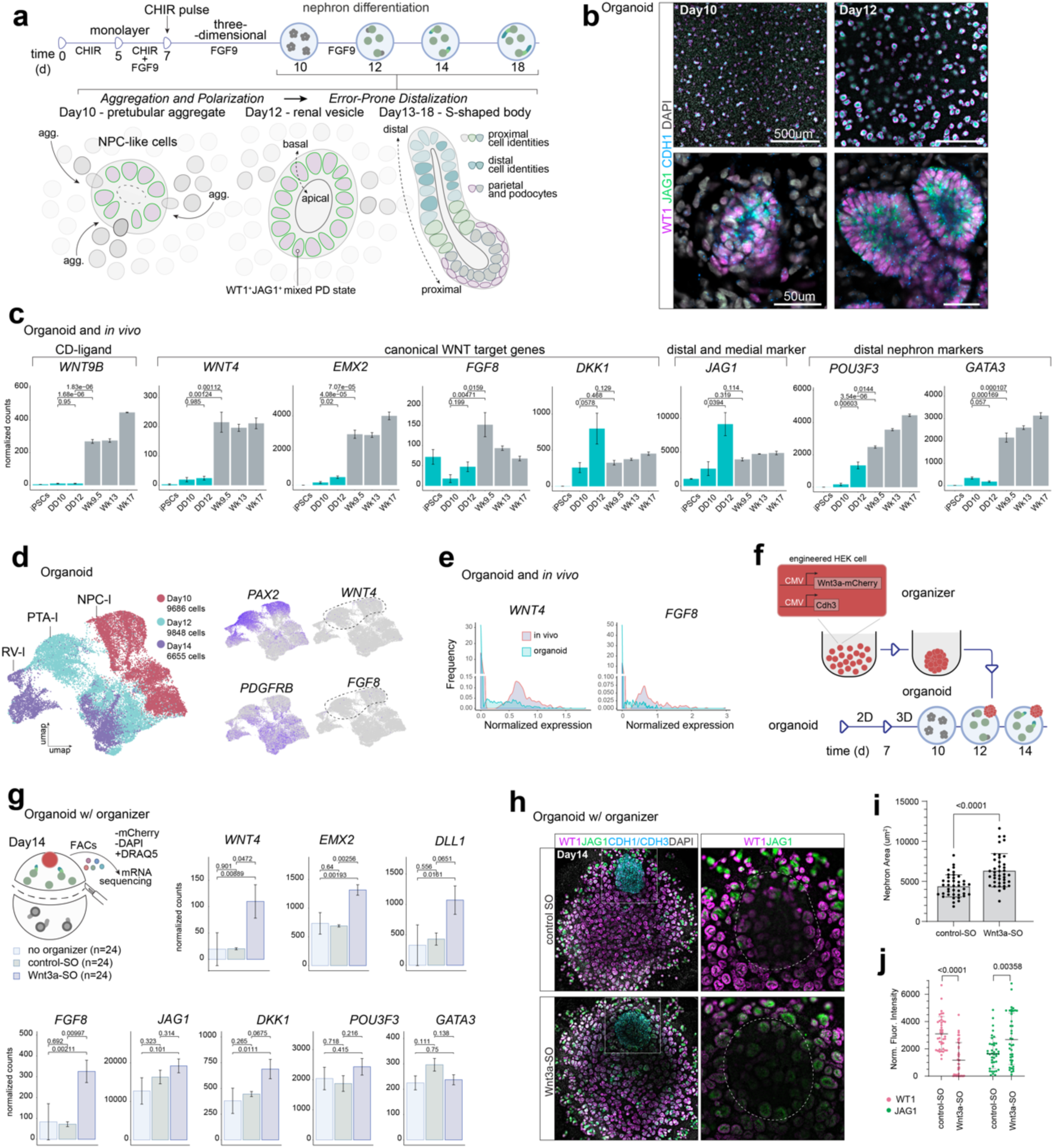
Kidney organoid nephrons form without intrinsic asymmetry but can respond to cell-secreted localized WNT ligands. **a,** Overview of kidney organoid differentiation protocol, depicting formation of nephron-like structures in the absence of CD, noting apical-basal polarity, PD-polarity, no AD-polarity. **b,** Organoid immunostains showing condensing PTA-like structures (Day10), lumenized epithelial RV-like structures (Day12); stains and scale bars as shown. **c,** Bulk transcript count comparisons between kidney organoids, developing organoids, and iPSCs. **d,** UMAP plots showing organoid scRNAseq for days 10, 12, and 14, with lineage and differentiation markers shown as feature plots. **e,** Histograms comparing number of cells with expression of *WNT4* and *FGF8* between developing kidney and kidney organoid single-cell transcriptomic datasets. **f,** Schema showing strategy for providing localized canonical WNT ligands to kidney organoids. **g,** Bulk transcript count comparisons of Day14 organizer-coupled organoid regions for controls, WNT-secreting organizers, and non-coupled organoids. **h,** Day14 organoids coupled with control or WNT secreting organizers immunostained as shown. Right panels show magnification of left panel inlets (white square); dotted lines demarcating organizers. **i-j,** Graphs showing nephron areas and quantified immunofluorescence.

Rosettes of WT1^+^ cyst-like structure of around 35-45 µm in diameter, reminiscent of PTAs, are visible at differentiation day 10 (**Fig.2b**), indicating that nephron precursors begin to epithelialize 3 days after pulsing with CHIR. The partially lumenized epithelial cell clusters are positive for Notch-ligand JAG1 and epithelial cadherin CDH1 and within 48 hours of forming (day 12), they become fully lumenized epithelia of around 55-65 µm diameter, morphologically reminiscent of RVs (**Fig.2b**). However, immunolabelling of proteins WT1, JAG1, and CDH1 indicate these structures display radial symmetry and there are no morphological cues of polarized or gradual recruitment of NPCs. This is contrary to what is observed in human kidney development *in utero*, where NPC recruitment is polarized towards the CD-distant point while *WT1* and *JAG1* expression diverges in the RV into proximal and distal-medial regions, respectively (compare **Fig.2b** to **Fig.1e** and **Fig.S2c**).

To assess whether organoids display evidence of canonical WNT signaling we compared their transcriptional profiles to human kidney development. We performed bulk RNA sequencing across day 10-12 organoids and compared these to a range of human kidneys at weeks 9, 13, and 17; data from ^11^. In organoids, except for *WNT3* other canonical WNTs (e.g., *WNT9B* and *WNT3A*) were expressed below reasonable transcript count threshold levels, as were nephron canonical WNT targets *WNT4* and *FGF8* (**Fig.2c; Supplementary table 2**). This is in contrast with kidneys, where *WNT9B* and its targets *WNT4* and *FGF8* were readily detected, as expected from a canonical WNT response being driven across the kidney cortex where active nephron formation is occurring.

The complete lack of a canonical WNT response is surprising given its known importance during *in vivo* nephrogenesis. To validate that organoid nephrons develop without triggering a canonical WNT signaling event, we scrutinized the organoid PTA to RV nephron transition when WNT signaling would be expected. Single cell RNA sequencing data capturing days 10, 12 and 14 of organoid differentiation show that a PTA-like cell state is present (low *MEOX1*, *PCDH15, CITED1,* and positive for *EMX2*, *JAG1*, *LHX1*, *KRT8*), however, it resembles a hybrid NPC-PTA cell state with undefined boundaries between NPC and PTA profiles (compare **Fig.2d Fig.S4a-b** with *in vivo* gene expression plots Fig.1i for *MEOX1*). During the developmental range from PTA to RV, *WNT4* and *FGF8* mRNA transcripts were absent or very low in the organoid nephron while they are readily detected in *in vivo* samples capturing the equivalent range of developmental stages (**Fig.2d,e**: data analyzed from our recent work ^37^ and *in vivo* data as shown in Fig.1). The organoid NPC and PTA also display unusual expression profiles of mixed nephron and interstitial characteristics. *In vivo*, *ELAV4* is restricted to only the nephrogenic lineage, but a *PAX2*^−^ (nephron lineage marker), *PDGFRB^+^* (interstitial marker), *ELVAL4^+^*cell is observed in organoids (**Fig.S4b**), thus representing a state not seen *in vivo*. Organoids did, however, express WNT receptors and signaling transduction modifiers from the Frizzled, Low-density lipoprotein receptor-related protein receptors, and R-spondin family members (*FZD1*, *FZD2*, *FZD4*, *FZD6*, *FZD7*, *FZD8*, *FZD9*, *FZD10*, *LRP5*, *LRP6, RSPO3*, *RSPO4* - notably not *RSPO1* or *RSPO2*) **Fig.S4c**. Notably, organoid NPCs expressed canonical WNT receptors FZD4, FZD7, and FZD5 in a PTA/RV-like subset, along with LRP5/6 and RSPO3-4, but not RSPO1-2 (**Fig.S4C**). This suggests that organoid nephrons are ready to respond to canonical WNT ligands such as WNT9B (FZD4, FZD5) or WNT3A (FZD4, FZD5, FZD7) ^38,39^ but do not do so because of the absence of the WNT secreting CD. There are differences in organoid and kidney expression profiles for non-canonical WNTs, and we readily detected *WNT11*, *WNT5A*, and *WNT5B* suggesting organoids only partially recapitulate the *in vivo* kidney state (**Fig.1c Fig.S4c** and **Supplemental table 2**).

*In vivo*, expression of *JAG1* and *EMX2* are thought to be downstream of the CD-secreted canonical WNT signal ^8,40–42^. In organoids, even in the absence of a canonical WNT, *JAG1* and *EMX2* transcripts rose ~ 3-fold between days 10 and 12 (**Fig.2b,c S4b**). On day 14, genes marking distal nephron cells (e.g. *POU3F3*) were detected, but a lack of *GATA3* transcripts indicates minimal specification of a *bona fide* connecting tubule (CNT), a key distal nephron derivative (**Fig.S4b**).

Our findings highlight a model where kidney organoids contain NPC-like cells that undergo differentiation, epithelialization, and upregulation of *JAG1* in the absence of polarizing ureteric stalk-derived *WNT9B*, leading to RV-like structures devoid of *WNT4 and FGF8* expression and polarity.

### Canonical WNT-secreting signaling centers drive β-catenin target genes in organoid nephrons

Given that nephron cells that emerge in organoids possess canonical WNT pathway components, we hypothesized that their lack of WNT target gene activation is to be imputed to a lack of canonical WNT ligands. To test this hypothesis, we used WNT-secreting synthetic organizer (SO) cell clusters. These SOs are composed of HEK cells that are genetically engineered to produce either a control fluorescent protein or Wnt3a in a constitutive (Wnt3a-SO) manner. All SO cells express *Cdh3* for increased self-aggregation and mCherry as fluorescence marker (**Fig. 2f**) ^32^. Wnt3a-SOs and only overexpress *Wnt3a* and not other canonical Wnt ligands, while control-SOs do not express any canonical Wnts; as confirmed by bulk RNA sequencing on the organizer cells themselves (**Supplemental table 4**). To confirm that these cell clusters can elicit nephrogenic and canonical WNT responses we tested their action in intact nephron-forming niches in embryonic mouse kidney explants. Small (5,000 cells) Wnt3a-SOs and control SOs were assembled in ultra-low adhesion wells for 24h and positioned adjacent to embryonic explanted kidneys. As shown in **Fig. S4d,e**^32^, ectopic Jag1^+^/Cdh1^+^ nephrons formed in response to Wnt3a-SO but not in SO controls, confirming that Wnt ligands produced by synthetic organizer cells can induce a canonical Wnt-response in developing mouse kidney NPCs. To determine if nephron polarization can be manipulated via synthetic organizers in human iPSC-derived nephrons, we introduced Wnt3A-SO and control SO on day 12 organoids, when nephrons are non-polarized RV-like structures. We transcriptionally profiled those nephrons closest to the SOs after 2 days of signaling at differentiation day 14 (**Fig.2f,g**) and enriched for organoid cells closest to the organizers by mechanically dissecting the region of cells in proximity to the organizers and removed SO cells based on negative FACS sorting for mCherry. Exposure to Wnt3a-SOs increased expression of *EMX2*, *FGF8,* and *DLL1* (**Fig. 2g**). *WNT4* transcript counts remained at subthreshold levels with a trend increase to Wnt3A-SO, likely due to *WNT4* being expressed at low levels in general and the diluting effect of other interstitial cell populations in the organoid. Of note, β-catenin target gene *DKK1* was increased in response to Wnt3a but neither *POU3F3* nor *GATA3* (distal patterning genes) responded to the ectopic Wnt3a ligands within the 48-hour timeframe. This is consistent with the *in vivo* setting where *EMX2* and *FGF8* are expressed prior to *POU3F3* and *GATA3* adjacent to the*WNT11/WNT9B* transition point (compare to Fig.1c).

To assess how individual nephrons responded to the introduction of ectopic Wnts we assessed markers for the early CD-adjacent (JAG1^+^) and podocyte differentiation cell states (WT1^+^) alongside epithelial marker CDH1 in organoids conjugated with control and Wnt3a-SO. As shown in **Fig. 2h**, nephrons close to control SOs are unaffected compared to neighboring structures, but in Wnt3a-SO conjugated organoids, nephrons close to Wnt3A-SOs grew larger and downregulated WT1 while upregulating JAG1 (**Fig.2i,j**). This is consistent with our previous work detailing β-catenin acting as an antagonist to proximal and podocyte development in mice ^20^. We noticed that the patterning effects were local and affected only nephrons in the vicinity of the organizer and not regions of the organoid farther from the organizer. To quantify the range to which β-catenin signaling is affected in the organoid, we assessed LEF1 (β-catenin target) protein levels and observed increased LEF1 protein abundance up to hundreds of microns away from the organizer cells (**Fig.S4f,g**).

These results demonstrate that engineered ectopic synthetic cellular organizers can express a canonical Wnt ligand that drives β-catenin mediated responses in human kidney organoid cells, locally program nephrons to upregulate early distal cell markers (*EMX2*, *FGF8*) and introduce developmental programs otherwise missing in hiPSC-derived organoid cultures.

### Tunable WNT-secreting synthetic organizers drive nephron axial symmetry-breaking and morphogenesis

Building on these results, we engineered HEK organizer cells with tunable control of canonical Wnt ligands *Wnt3a* and *WNT9B*, reasoning that adjustable control of the production of Wnt ligands from organizer cells could allow us to regulate patterning in the organoids. To do so, we generated new organizers producing Wnt/WNT ligands (Wnt3a and WNT9B) under a dox-inducible promoter, alongside constitutive expression of cadherin protein 3 (*CDH3*) (**Fig. S5d**) ^43,44^. Inducible WNT SO are from here on referred to as iWnt3a-SO and iWNT9B-SO. Flow cytometry analyses showed an expected sigmoid dose-response curve of inducible expression of *TagBFP-2A-Wnt3a* transgene from iWnt3a-SO cells (EC_50_ of 2.14±0.74nM), peaking past 10nM (**Fig.S5a-c**), supporting capacity of the system to express WNT transgenes in graded manner. To test the capacity of these iWnt-SO cells to induce WNT/β-catenin transcriptional responses in neighboring cells, we set up co-cultures of iWnt-SO cells with mouse ES reporter cells carrying a β-catenin TCF/LEF-GFP reporter ^45^. When induced with doxycycline to express the WNT ligand, iWnt3a cells drove reporter activation in neighboring mES cells (**Fig.3a**, **Fig.S5e**). Similar experiments with iWNT9B-SO cells, initially gave no activation of the reporter ES cells. Given that the co-receptor RSPO1 is a critical potentiator of WNT9B-FZD5 interactions and downstream β-catenin activity ^46^ we tested if adding RSPO1-conditioned media in co-culture experiments with iWNT9B can sensitize TCF/LEF mESC reporters to WNT9B. Together, RSPO-1 and WNT9B drove robust ES cell reporter activation, demonstrating capacity of iWNT9B cells to induce WNT/β-catenin responses in neighboring cells in presence of RSPO-1 (**Fig.3b** and **S5f**). Given the difference between the Wnt3a and WNT9B results, we further tested the capacity of iWNT9B-SOs to induce tunable nephrogenesis in mouse kidney explants. To do so, we conjugated iWNT9B-SOs to mouse kidney explants in the presence of increasing doses of dox. iWNT9-SOs drove differentiation of Jag1^+^ nephron precursors adjacent to the organizers in a dose-dependent fashion (**Fig.S5g**). Of note, WNT9B did not require external supplementation of RSPO-1 in kidney explants. The *Rspo1* co-receptor is highly expressed in NPCs and early nephrogenesis *in vivo*; single cell RNA sequencing from mouse nephrogenesis ^47^, consistent with conserved expression in human (Fig.1k).

**Figure 3.**
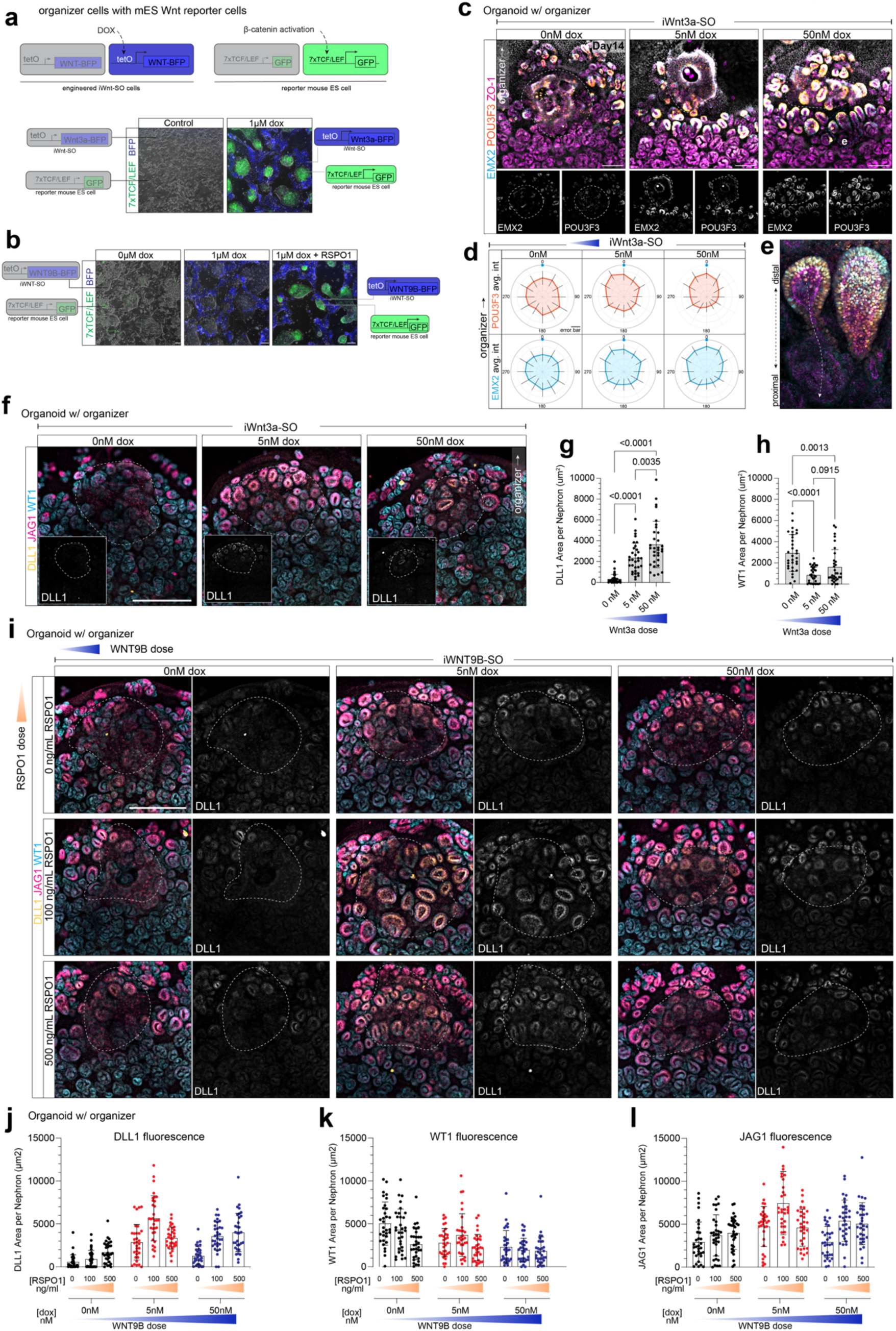
Organoid nephrons distalize and align their nascent axial polarities towards tunable WNT-secreting organizers. **a,** Overview of doxycycline-inducible WNT synthetic organizers that report WNT transgene expression via TagBFP (iWnt-SO), and receiver mouse embryonic stem cells that report β-catenin activity via eGFP. **b,** Live fluorescence microscopy of co-cultures of iWNT-SO and embryonic stem cells reporters in the absence or presence of dox, for iWnt3a-SO cells (top) and iWNT9B-SO cells (bottom); iWNT9B-SO cells with or without RSPO-1. **c,** Day14 organoids coupled with iWnt3a-SOs at different doxycycline dosages, immunostained as shown. **d,** Radar plots showing distribution of POU3F3 and EMX2 fluorescence in nephrons relative to the nephron lumen (radar center) and organizer-adjacent point (blue dot) for nephrons shown in (c). **e,** Confocal close-up of iWnt3a-SO polarized nephrons (from blue dashed area in (c)), stained as show and with noted proximal-distal axis. **f,** Organoids treated as in (c) immunostained for as shown. **g-h,** Graphs showing quantified nephron areas positive for DLL1 (g) and WT1 (h) per nephron as shown in (f). **i,** Day14 organoids coupled with iWNT9B-SOs at different doxycycline dosages and over increasing RSPO1 dosages, immunostained against as shown. **j-l,** Graphs showing quantified nephron areas positive for DLL1 (j), JAG1 (k), and WT1 (l) per nephron as shown in (i).

With these validated tunable iWnt-SO, we tested if they could drive nephron patterning in human kidney organoids. To do so, we coupled kidney organoids with iWnt-SOs at day 12 when organoid nephrons begin to polarize along their PD axes. These organizer-to-organoid coupled tissues were cultured in media with varying amounts of dox, thus inducing increasing levels of Wnt ligands from the organizer cells. After 48h of culture in these conditions, we profiled nephron axial polarities by immunofluorescent stains for nephron patterning markers POU3F3 (distal), EMX2 (distal transcription factor), JAG1 (distal/medial), DLL1 (medial/proximal Notch ligand), and WT1 (proximal). EMX2 and DLL1 were selected as are highly sensitive markers for cells responding to constitutive Wnt3a (Fig.2g). iWnt3a-SOs drove a dosage-dependent upregulation of WNT/β-catenin and nephron distal patterning markers EMX2 and POU3F3 and induced morphological changes (**Fig.3c**). To define patterning and morphogenetic responses, we assayed the distribution of EMX2 and POU3F3 by measuring the polarization of markers within individual nephron tubules relative to the position of the organizer (similarly to how we established *in vivo* nephron axial asymmetry in **Fig.1**). The position of these distal markers was enriched in the part of the nephron closest to the organizer. Moreover, and the number of nephrons that polarized towards the Wnt3a source increased (**Fig.3d-e**, **Fig.S5g-j**). In addition, DLL1 was similarly upregulated in a dosage dependent manner, whereas proximal marker WT1 was downregulated (**Fig.3f-h**). Intriguingly, nephron morphogenesis was affected, as nephrons elongated towards the Wnt3a source, and their PD length was increased compared to controls (**Fig.S5h**).

We next determined whether WNT9B-producing organizers could generate similar changes in organoids. To do so, given the requirement for RSPO-1 to enable iWNT9B organizers to drive a canonical signaling response in adjacent cells, we built a WNT9B/RSPO-1 matrix of culture condition to expose organoid nephrons to varying dosages of each. As shown in **Fig.3i-l**, and similarly to iWnt3a results, overall presence of both RSPO-1 and WNT9B resulted in upregulation of DLL1 and JAG1, and downregulation of proximal marker WT1. In these experiments, we noticed that nephrons were highly sensitive to changes in WNT9B and RSPO-1: DLL1 is virtually absent in organoids where *WNT9B* is not induced (0nM dox column) but becomes strongly upregulated in 5nM dox *WNT9B* induction at any RSPO-1 dosage. A higher DLL1 increase was observed at 5nM dox with 100ng/mL RSPO-1 compared to the 50nM dox equivalent, an effect lost at the 500ng/mL RSPO-1 dose. This suggests that nephrons in human kidney organoids respond in dosage-dependent manners to WNT ligand inputs (**Fig.3i-j**; statistical analyses in **Fig.S5l-m**).

These results show that iWnt-SOs can drive dose-dependent WNT/β-catenin signaling in iPSC-derived nephrons leading to nephrons aligning and patterning their PD axes towards the synthetic organizers.

### WNT-secreting synthetic organizers tunably orchestrate nephron patterning and morphogenesis

We have shown that the initial events of patterning and morphogenesis of nephron development can be controlled by WNT-producing synthetic organizers. We therefore asked if continuous exposure to Wnt ligands, as is observed *in vivo* (see **Fig.1**), affects further nephron patterning and morphogenesis *in vitro*.

To explore this, we conjugated synthetic organizers constitutively expressing *Wnt3a* with kidney organoids at day 12 and analyzed patterning outcomes at day 18. To assess nephron patterning, we labeled epithelial cells of the distal nephron by immunostaining for CDH1 and POU3F3 and similarly marked apical surfaces of proximal renal corpuscle cells (podocyte and parietal epithelium) as well as the apical surface throughout the epithelial component of the nephron by labeling tight junction protein ZO-1 ^16^. In control SO-conjugated organoids, nephrons positioned under organizers developed normal renal corpuscles (strong apical ZO-1), and short distal tubules (POU3F3^+^), similar to other regions of the organoids distant from the organizer. However, exposure to Wnt3a-SOs transformed nephrons adjacent and under the organizer into distalized POU3F3^+^ tubules devoid of renal corpuscles-like structures (**Fig.4a** and **Fig.S6a**). To assess these effects in detail, we defined the SO-adjacent nephrons according to their distance to the organizer and their position within the organoid, and their patterning. Their positions were classified into an “under” region that is underneath the organizer, an “in” region this is from the organizer towards the center of the organoid, and an “out” region that is equivalent in dimension to the “in” region but instead accounts for nephrons positioned towards the periphery of the organoid (each marked on **Fig.4a**). Nephron patterning were categorized as large POU3F3^+^ nephrons, small truncated POU3F3^+^ cell clusters, and strongly positive ZO-1 renal corpuscles.

**Figure 4.**
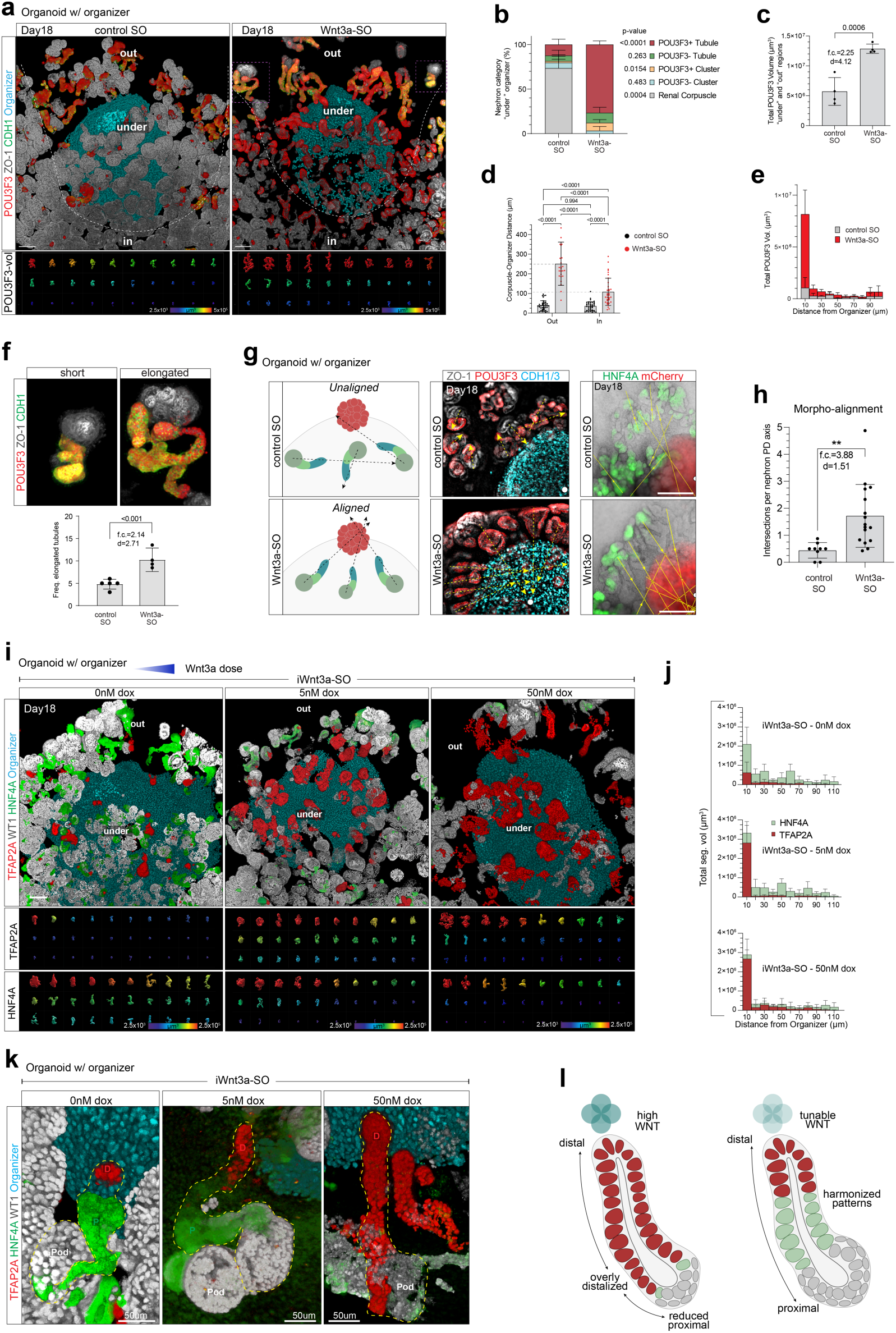
Organoid nephron proximal-distal patterning and polarization can be controlled by Wnt-secreting synthetic organizers. **a,** 3D renders of Day18 organoids coupled with control spheroids or Wnt3a organizers, immunostained for distal tubule markers POU3F3 and CDH1, and the tight-junction marker ZO-1. Bottom panels show top 30 POU3F3 objects descending by volume, for nephrons located under the organizer and towards the organoid’s periphery (under-and-out, above dotted line). **b,** Percentages of nephron categories underneath organizers shown in (a). **c,** Total POU3F3 volume from nephrons located in the under-and-out position. **d,** Distances between organizer peripheries and nearest renal corpuscles located towards the organoid’s periphery (out) or center (in). **e,** Breakdown of total POU3F3 volumes from (c), showing POU3F3 volume distributions relative to the distance from the organizer. **f,** Number of nephrons showing large POU3F3 volumes per organoid. **g,** Tubule morpho-alignment concept (left), immunostained examples (middle), and fluorescent reporters (right) highlighting alignment of nephron PD axes towards the Wnt3a sources. White circle marks the organizer center; yellow arrows mark tubule trajectories **h,** Morphoalignment measurement of number of tubule path intersections relative to the number of paths drawn per organoid. **i,** 3D renders of TFAP2A, HNF4A, and WT1 structures in iWnt3a-SOs coupled organoids in Day18 organoids at different doxycycline dosages. Bottom panels show top 30 POU3F3 and HNF4A objects descending by volume, for nephrons located under the organizer and towards the organoid’s periphery. **j,** Distribution of TFAP2A (top panel) and HNF4A (bottom panel) volumes relative to the distance from the organizer for nephrons in the under-and-out position. **k,** Close-ups of nephrons surrounding iWnt3a-SOs showing tubular alignment and patterning at basal, intermediate, and peak dosages. **l,** Proposed model: Constitutive provision of canonical WNT/β-catenin ligands at high levels distalizes developing nephrons at the expense of proximal identities. Tuning WNT levels induces distalization and morphoalignment, while minimizing suppression of proximal segments, yielding nephrons with continuous patterning.

These patterning analyses showed that nephrons forming under the Wnt3a-SO did not develop renal corpuscles and instead generated POU3F3^+^/CDH1^+^ tubules, contrary to control-SO conjugated nephrons that develop normal renal corpuscles in that same region (**Fig.4b**). This absence of renal corpuscle (podocyte and parietal epithelium) cell fates in response to Wnt3a together with the distance-dependent abundance of LEF1 (**Fig.S4f**) suggest nephrons exhibit a Wnt3a dosage response that is mapped out differently at different distances to the organizer. Scrutinizing the spatial distribution of renal corpuscles relative to the organizer showed that in Wnt3a-SO conjugated organoids, nephrons developed renal corpuscles 108μm from the organizer, as measured on the “inside” region, while this distance increased to 251μm on the “outside” region (**Fig.4d**). In control SO-conjugated organoids, these distances were considerably shorter at 35.7μm and 39.1μm, respectively. Nephrons in the “under” and “out” regions of Wnt3a-SO conjugated organoids were more susceptible to this effect (**Fig.4d**). In the presence of Wnt3a, renal corpuscle cell fates were completely suppressed within a 100um radius of the Wnt3a-SO on the outside region. Conversely, distal POU3F3 segments were larger and distributed towards Wnt3a-SOs compared to controls-SO conjugated organoids (**Fig.4e**).

To confirm these axial patterning changes, we performed equivalent spatial distribution analyses for a range of PD cell markers: proximal (HNF4A^+^) and distal (TFAP2A^+^) nephron segments and marked podocytes (WT1^+^) for reference (**Fig.S6a-e**). Similar to our POU3F3/ZO-1 analyses, large distal TFAP2A^+^ segments were enriched adjacent to Wnt3a-SO compared to control-SO (**Fig.S6b,c**), while HNF4A^+^ segments objects were depleted and smaller; no HNF4A^+^ segments above 2×10^3^ μm^3^ were found within a 70-80μm range from Wnt-sources while being abundant in controls (**Fig.S6c**). These data demonstrated a patterning effect on nephrons that is dependent on the distance from the organizers.

To corroborate the nephron patterning changes driven by Wnt3a, we performed spatially restricted single cell RNA sequencing (scRNA-seq) from Wnt3a-SO and control-SO coupled organoids. Following cell clustering analyses, cell groups reflecting distal (*POU3F3*), proximal (*HNF4A*), and podocytes (*NPHS1*) identities were identified (**Fig.S7a-g**). As expected, transcript reads for WNT/β-catenin-target genes known to be expressed in nephrogenesis (*e.g., LEF1, AXIN2, DKK1, HNF1B, FGF8*) were more frequently detected in nephron cells next to Wnt3a-SOs than control-SOs. Markers for distal tubules (*LGR5*, *POU3F3, TFAP2A*), loop-of-Henle and macula densa (*IRX1*, *PAPPA2, SLC12A1, FGF8*), and connecting tubules *(GATA3*) were similarly enriched in Wnt3a-SO samples, whereas proximal (*HNF4A*) and podocyte markers (*WT1, MAFB, PODXL*) were minimally downregulated (**Fig.S7h**). The relatively high abundance of podocyte and proximal markers reflects the short distance that these cell identities were suppressed on the “*inside*” organoid facing surface (**Fig.4a**, **Fig.S7a**).

Nephrons developing adjacent to Wnt-secreting SOs displayed distinct morphological changes. To better understand these changes, we analyzed nephron morphologies by generating a collection of 3D-rendered nephrons. As shown in **Fig.4a** (bottom row) and **Fig.4f** (top row), distal nephron tubules appeared larger in Wnt3a-conjugated nephrons compared to control-SO nephrons. To quantify this, we calculated the volume of POU3F3^+^ nephron domains developing in the “out” and “under” region of Wnt3a-SO conjugated organoids. These displayed a 2.26-fold increased volume of in Wnt3a-SOs compared to controls (**Fig.4c**). Some nephrons in Wnt3a-SO conditions developed a particularly interesting phenotype, reminiscent of elongating distal tubule structures observed *in vivo*. Categorizing these as ‘*elongated*’ in contrast to the shorter nephron ‘*short*’, we defined the elongated phenotype within a volumetric bracket (see methods). Elongated nephron structures were found at a 2.14-fold increased abundance near Wnt3a-SO compared to control organizers (**Fig.4f**).

Given that nephrons align towards the WNT11/WNT9B transition point *in vivo*, and they start to do so in our tunable iWnt-SOs (Fig.3), we characterized the alignment of the nephron structures towards the organizers in these day 18 conjugated organoids. To do so, we mapped the orientation of tubules relative to the organizer by defining nephron PD axial trajectories with respect to the organizers. We determined tubule orientations using PD markers (using immunolabelling and fluorescent reporter iPSC lines) and quantified the convergence of PD axes (**Fig.4g-h**, and **S8a**). The projected paths of nephron tubules converged more frequently and closer to Wnt3a-SOs compared to controls-SOs, demonstrating that nephron tubules aligned their PD axes towards the WNT source.

These data show that nephron morphogenesis and differentiation are affected by Wnt3a-SOs such that organoid nephrons orient their distal tubules towards the Wnt3a-SO and develop distal nephron cell identities in a distance-dependent manner that simultaneously suppresses proximal tubule and renal corpuscle cell fates.

Exposure to high levels of Wnt3a expanded distal segments at the expense of proximal identities (highlighted for example in Fig. 4b), thus generating a non-physiologic pattern of gene expression in nephrons. To determine if tuned Wnt3a sources can evenly pattern nephrons without suppressing proximal and podocyte identities, we evaluated TFAP2A (distal), HNF4A (proximal) and WT1 (podocyte) nephron segment sizes in day 18 organoids coupled with iWNT3a-SOs at different Wnt/dox dosages. 3D reconstruction of confocally scanned nephrons show normal WT1^+^ renal corpuscles and elongated HNF4A^+^ proximal precursor segments underneath iWNT3a-SO in the absence of doxycycline (**Fig.4i**).

At intermediate Wnt dosages (5nM), continuity of TFAP2A-HNF4A-WT1 segments could be seen in nephrons adjacent to the organizer and towards the organoid’s periphery. Prominent TFAP2A^+^ distal nephron segments formed underneath the organizer in the absence of elongated HNF4A^+^ segments and renal corpuscles developed, albeit smaller. At peak Wnt dosages (50nM), large TFAP2A^+^ structures similarly appeared underneath organizers and WT1 expression was partially suppressed and HNF4A^+^ tubules were excluded to positions further from the organizer. To quantify these observations, we rendered and measured TFAP2A and HNF4A segment volumes in relationship to their spatial distribution. Distal segments (TFAP2A^+^) increased in volume at increasing Wnt doses while correspondingly, HNF4A segment volumes decreased at the highest dose of Wnt (**Fig.S8b,c**). At high Wnt3a doses the tunable system mirrored the distalization and suppression of proximal identities observed earlier (**Fig.4a**). Strikingly, at intermediate Wnt3a doses, both TFAP2A^+^ and HNF4A^+^ segments formed that connected within single nephron tubules (**Fig.4k**). This was reflected by measurements of the size distribution of segment volumes. Only in intermediate Wnt3a doses did HNF4A^+^ and large POU3F3^+^ segments form immediately adjacent to the organizer (**Fig. 4j**), Similarly, at intermediate Wnt3a doses renal corpuscles formed adjacent to the organizer but their development was suppressed by high Wnt3a doses (**Fig.S8d**). This is consistent with distal differentiation being driven near the Wnt source and at a high dose and suggests that intermediate Wnt doses are compatible with a fully patterned nephron (**Fig.4i-k**).

Collectively, these experiments demonstrate that canonical Wnt ligands can, when provided at the right levels, drive development of well-patterned nephron segments and polarize nephron axial morphogenesis towards the Wnt source (**Fig.4l**).

## Discussion

### Spatial transcriptional analyses highlight a new axial polarity during nephron development

Our findings describe an axial polarity detailing nephron cells’ relationship to the CD. This is configured as NPCs positioned around the CD tip undergo a mesenchymal to epithelial transition to generate nephrons and thereafter the axis persists through the stereotyped developmental morphogenetic steps of the PTA, RV, CSB, and SSB impacting adult nephron morphologies and function. Our spatial transcriptional analyses show that a CD *adjacent-distant* (AD) polarity precedes the formation of the PD polarity. The AD polarity initially traverses the partially epithelialized PTA and can be defined in the PTA by how cells are positioned in relation to the WNT11/WNT9B transition point and the WNT9B^+^ CD surface (shown in Fig.1). Subsequent to PTA stages, the CD-adjacent domains define the nephron epithelium that is adjacent to the WNT9B^+^ CD stalk as the RV and CSB grows and elongate and the SSB folds into its typical structure. Adding the AD axis as a reference point to describe nephrogenesis assists us in defining gene expression patterns and areas of the forming nephron and it integrates with the proposed gradual recruitment model and a time-dependent mechanism dictating nephron fate specification through sequential activation of signaling pathways ^15^.

Providing a new term to an existing language model will create discussion on whether current nomenclature suffice. We anticipate that the AD axis will be welcomed as the kidney organoid field tackles questions on how to replicate stereotyped developmental steps *in vitro*. Further, it assists with changing misconceptions about the PTA. The PTA has long been described as entirely non-epithelial, but its nature is nuanced and consists of partially epithelial cells in CD adjacent positions that epithelialize and asymmetrically upregulate gene expression while new NPC recruits are slotted into the forming nephron; a process intrinsically linked to cell recruitment and gradual formation of spatial positional identities. Given that this asymmetry appears to play a crucial role in nephron patterning and therefore acquisition of physiologies, we find it to be necessary to consider.

Is the AD axis temporary during nephrogenesis? While the precursor-progeny relationships of AD axial positions need to be defined in future experiments, our data shows that molecularly distinct AD regions are prominent in the RV, CSB, and SSB, thus making it hard to envisage how SSB and subsequent nephron patterning are not influenced by this polarity. Putative precursors can be defined by it. For instance, in the SSB we detect a secondary upregulation of *WNT4* in CD-adjacent proximal precursors; an expected find as we have previously shown *in situ* hybridization of this domain ^48^. However, the dynamics of WNT4 illustrates the asymmetry of the nephron and puts forward an explanation for why podocytes are mosaically labeled with *Wnt4^Cre^* fate mapping ^48,49^. Low expression of WNT4 in CD-distant cells is a likely reason for previously observed incomplete *Wnt4 ^Cre^* mediate recombination in CD-distant podocytes, while other nephron segments are robustly labeled with Cre-dependent reporters ^48^. Moreover, an AD axis also begins to shed light on the chirality of SSBs, as one axis is insufficient, but two axes can define chiral structures ^16^.

The perhaps most obvious reason for why an AD axis is required can be demonstrated by a simple visualization. Picture the SSB, rotate it on its AD axis, and imagine how it now connects with the CD and forms podocytes in close contact with a developing macula densa - a hard feat. We now need to define AD fate-maps, better understand AD spatial transcriptional signatures in three dimensions, and determine the downstream functional significance of this interesting axis.

### Building an artificial kidney

A cell derived kidney suitable for replacement therapies should be connected to a vascular in and output, contain a physical filter, perform solute reabsorption, and lead unwanted solutes and metabolites to a drainage system. The latter function is normally performed by a luminal connection between the distal end of the nephron and the CD. Given that this is a critical requirement for function, we therefore sought to delineate how the distal nephron is polarized towards the CD and replicate this in an organoid model with a view to develop an approach for channeling urine in an artificial kidney.

To control nephron patterning in hiPSC kidney organoids we introduced WNT-secreting cellular organizers to replace CD-derived WNT9B ligands. This drove distalization and physical nephron alignment towards the WNT source. The concept of organizers has been one of the most powerful developmental biology concepts since the Spemann/Mangold organizer experiments. Although replicating sources of morphogenetic factors with beads and ectopic tissue has been performed before *in vivo* and in organoids ^33,50^, it is only recently that there have been attempts to generate synthetic organizers made of cells. In these studies, organizing cells are placed in the context of a stem-cell derived construct, where they instruct polarization and cell fate changes. In our study, we utilized self-organizing HEK clusters that overexpress cadherin, possibly contributing to their compactness and capacity to stay coherent when integrated in the organoids. In fact, using hiPSC organizer cells resulted in integration of the hiPSC ‘organizer’ cells in the rest of the organoid, generating a chimeric structure ^33^. Here, the organizer cells do not lose compaction and stay a separate entity compared to the organoid. This allows the organoid cells to continue their developmental trajectory without being invaded by organizer cells. In the later days of the organoid/organizer co-cultures, we could observe that the nephron tubules extend towards the organizers, and burrow into the organizer itself, as if they were reaching out to connect to a structure in an attempt to make epithelial connections. In future experiments it will be important to develop organizer cells of an CD-like epithelial nature, to engineer a continuous lumen between CD and nephron.

In our hands, providing organoid nephrons with WNT secreting organizers distalized nephrons. This demonstrates that human iPSC derived nephrons respond to canonical WNTs by upregulating a transcriptional response akin to that expected *in vivo*. However, it is surprising that we found no evidence for an endogenous canonical WNT ligand-receptor pairing in control organoids during the timeframe when nephrons form and pattern. As part of the standard differentiation protocol ^21^ cells are pulsed with GSK3β-inhibitor CHIR at day 7 and nephrons emerge 3 days later at day 10. There is therefore a discrepancy between when β-catenin signaling is driven and when nephrons form, and the timing disagrees with a proposed *in vivo* developmental timeline for the PTA-RV progression ^11^. *In vitro* during days 10-14, expression of *WNT4* and *FGF8* (thought to be direct β-catenin targets) was low by bulk RNA sequencing (samples sequenced to ~20M reads) and by single cell RNA sequencing (~2500-4000 genes per cell). We therefore considered neither to be actively expressed. How then do nephrons form? In mice, ectopic Notch signaling can compensate in Wnt-deficient backgrounds when a dominant active Notch intracellular domain is overexpressed ^51^. *In vivo* experiments in mice have also shown that the nephrogenesis and activation of *Wnt4* and *Lhx1* require expression of *Fgf8* ^52^. In metanephric mesenchyme knockouts of *Fgf8*, NPCs aggregate into PTAs and RVs but lack *Wnt4* and *Lhx1* expression indicating Fgf8 is required during canonical Wnt activation or prefigures NPCs to be response. In recent work removing *Fgf8* in the NPC population, the NPCs are loosely packed but consistent with the full metanephric mesenchyme knockouts, removing *Fgf8* using either Pax8 or Wnt4 Cre lines shows PTAs still form ^53^. This agrees with our *in vivo* human results where FGF8 is present in the NPC population and is enriched in the CD-adjacent cells and *FGF8* expression precedes broad activation of WNT4 in the PTA, in CD-adjacent cells at the WNT11/WNT9B transition point. While this is therefore consistent with FGF8 contributing to cell aggregation it also demonstrates that it is unlikely to be an absolute requirement.

Since Notch is not directly manipulated here, *FGF8* is not detected using the methods we describe, and cells still aggregate and form PTAs, could they drive nephrogenesis through a Notch-dependent compensatory mechanism? Notch pathway and target genes (e.g., JAG1 and HES1) are expressed during the day 10-12 timeframe and delineating whether Notch can compensate for a lack of WNT input during organoid nephrogenesis should a focus of future studies that determine how to improve *in vitro* nephrogenesis and patterning. Of note, while we do not detect canonical WNT ligands in organoids, we did detect increased expression of WNT pathway modifiers *SFRP1*, *SFRP2*, *DKK1*, *DKK3* between days 10 and 12. These are genes often associated with β-catenin signaling and it remains possible that a basal level of canonical WNT signaling is occurring that we cannot detect with current methods. However, given that the organoids displayed expression of WNT receptors and co-receptors we favor the hypothesis that they are prefigured to receive a canonical WNT, do not normally do this due to the lack of ligand inputs, and are highly when ectopic WNTs are provided. These responses are possible to detect using the methods used in this study.

In addition to the transcription response, we also showed that organoid nephrons align their epithelial axes towards the WNT source. Activation of β-catenin in NPC culture settings *in vitro* has shown that this is sufficient to drive a cell aggregation program similar to that observed during NPC aggregation into PTAs ^41,42^. NPCs with activated β-catenin signaling become more motile and adhesive to other differentiating NPCs. *In vivo*, the NPCs move over distance to leave the NPC niche and join each forming nephron. As outlined above, the dynamics of *FGF8* expression together with phenotypes makes this an unlikely candidate for being the NPC chemoattractant. Our data raises the possibility that a canonical WNT acts as one of the attractive signals that coax NPCs to leave their niche and aggregate, but it does not explain how NPCs are recruited from the NPC niche into the proximal ends of forming RVs (where there is no or little WNT9B). NPC behavior is highly dynamic ^54,55^ and they are capable of extending long cellular processes from their positions within the NPC niche to forming nephrons; indicating they respond to a long-range chemotactic signal ^15^. The alignment of organoid nephrons to the WNT source suggests that WNT9B can act as a chemoattractive signal and simultaneously regulates a transcriptional distalizing program. From the perspective of engineering a cell-derived artificial kidney, the duality of one signal driving two processes would be advantageous as it can be used to drive distal nephrons towards defined positions in a common epithelial drainage system.

It is interesting to note as it could constitute an example for other attempts to “fix” organoid patterning, that rationale was lent to this synthetic organizer approach from our *in vivo* spatial transcriptional analyses. These analyses of human developing kidney show that β-catenin mediated transcription is polarized to those nephron cells adjacent to the *WNT9B* source. Human orthologs of previously proposed canonical WNT targets e.g., *Emx2*, *Fgf8*, *Jag1*, *Lhx1*, *Gata3*, *Wnt4*, and *Lef1* are expressed in that domain, and these *in vivo* human data agree with strong evidence for crosstalk between the CD and nephron derived from mouse studies. In mice, *Wnt9b* is produced in the CD and genetic studies show it is required in the nephron lineage, mediating NPC self-renewal, differentiation, and nephron formation, cell-fates, as well as for the maintenance of nephron planar cell polarity in maturing nephron tubules ^5,7,56^. In our past work we have shown that manipulation of the early mouse nephron with Wnt agonists or antagonists distalizes and proximalizes nephrons, respectively ^20^, which is in line with evidence from the chick mesonephros, where mesonephric tubules suppress podocyte development and reorienting nephron PD axes in response to Wnt3a expressed in the chick embryo ^57^. Further, these nephron patterning effects can at least partially be mimicked in hiPSC organoids through positive and negative regulation of β-catenin signaling ^50,58^. Our spatial transcriptional data therefore confirm that this WNT9B to nephron cross-lineage signal transduction is likely in the human nephrogenic niche and overcome past limitations of predictions made from single cell RNA sequencing data where spatial data has been missing. This indicates that this pipeline of identifying polarizing activities in human development that are missing in organoids, and reconstituting them in organoids with synthetic organizers, is a viable strategy to correct organoids patterning.

Here we propose that nephrons possess a previously undefined axial polarity, show its relationship to a cell organizer in vivo, and demonstrate that we can reconstitute these patterning and morphogenetic events in stem cell derived nephrons via synthetic cell organizers. We expect these finds and new approaches to provide a framework for understanding human nephron development and to provide novel strategies for engineering an artificial kidney that drains urine to a common epithelium.

## Acknowledgments

We thank all past and present members of the Lindström Laboratory and express our gratitude to Dr. Andrew McMahon and Dr. Zhongwei Li for their input. We than Dr. Jamie Davies for gifts of plasmids and cells. We further thank Dr. Seth Ruffins and the Optical Imaging Facility for assistance with microscopy and Dr. Bernadette Masinsin, Dr. Jorge Contreras and the Flow Cytometry Facility for help with cell isolation. We thank Kirsten Frieda, Linus Eng, Jay Mehta and Elizabeth Collins for assistance with spatial transcriptomic work.

This work was funded by Lindström lab funding from USC Department of Stem Cell Biology and Regenerative Medicine Startup Fund (N.O.L), National Institutes of Health under award number NIH/NIDDK R01DK136802 (N.O.L), American Society of Nephrology/United States Department of Health and Human Services KidneyX (N.O.L), CIRM training grant EDUC4-12756 (F.G.), T32 Training Grant in Development, Stem Cells, and Regenerative Medicine from NICHD (T32HD060549) (J.S.), and Morsut lab via NIGMS of the National Institutes of Health under award number R35GM138256 (L.M.); the National Science Foundation award number CBET-2145528 Faculty Early Career Development Program (L.M.); NSF RECODE from CBET-2034495 (L.M.); USC Department of Stem Cell Biology and Regenerative Medicine Startup Fund (L.M.); Wellcome Trust, Leap HOPE (L.M.), Viterbi Center for CIEBOrg (L.M.); grant 2023-332386 from the Chan Zuckerberg Initiative Donor Advised Fund, CZI DAF, an advised fund of the Silicon Valley Community Foundation (LM).

## Author contributions

FG, CF, LM, and NOL conceived and executed the project. LM and NOL directed the research and acquired funding. CF, FG, JS performed experiments. FG generated synthetic organizers, CF and JS generated organoids, FG and CF captured and analyzed data, RC prepared samples for spatial transcriptomics, CF and NL designed spatial transcriptomic experiments, NK and FDK analyzed spatial data, BG and MT performed data sampling, JS and MA performed organoid optimization, JS generated sequencing data and organoids, FG, CF, LM, and NOL prepared and assembled figures. All authors read and provided feedback on the manuscript.

## Materials and Methods

### Contact for Reagent and Resource Sharing

Further information and requests for resources and reagents should be directed to and will be fulfilled by the Lead Contacts, Nils O. Lindström and Leonardo Morsut.

### Animal Studies

Institutional Animal Care and Use Committees (IACUC) at the University of Southern California reviewed and approved all animal work as performed in this study (protocol# 21179). All work adhered to institutional guidelines. Timed matings were set up to recover embryos at the appropriate age (embryonic day 12.5), sex not known. C57/Bl6J were used. Kidney explants were cultured as previously described ^15,20^.

### Kidney samples

Kidney samples were collected under Institutional Review Board approved protocols (USC-HS-13-0399 and CHLA-14-2211) as for previous study ^37^. Following patient decision for termination, informed consent for donation of the products of conception for research purposes was obtained and samples collected without patient identifiers. The person obtaining informed consent was different than the physician performing the termination procedure, and the decision to donate tissue did not impact the method of termination. Developmental age was determined according to the American College of Obstetrics and Gynecology guidelines. The kidney samples were 13 and 18 weeks of gestation with no sex reported.

### Cell culture

Human iPSCs were grown on matrigel-coated (Corning: 354277) 6-well plates in E8 medium (Gibco: A1517001). Kidney organoids were derived using a previously published differentiation protocol ^21,37^. HEK-293 cells were cultured in Dulbecco’s Modified Eagle Medium (DMEM; Gibco: 11965092) supplemented with 10% fetal bovine serum (FBS; Genesee Scientific: 25-514, lot P085877), 2mM L-glutamine (Gibco: A2916801), 100U/ml;100μg/ml penicillin/streptomycin (Gibco: 15140122) and 0.1mM β-mercaptoethanol (Sigma Aldrich: M3148-25ML). HEK-293 cultures were grown to 90% confluence and passaged every 3-4 days at a 1:10-1:12 ratio. HEK-293 cells were harvested by aspirating culture medium, washing with phosphate-buffered saline (PBS; Gibco: 10010049), incubating with 0.35% trypsin (Genesee Scientific: 25-510) at 37°C for 3-5 minutes, neutralizing with double the volume of fresh medium, and collecting the desired ratio for passaging and organizer production.

### Directed differentiation to generate kidney organoids

Kidney organoid differentiation was adapted from published protocols ^21,59^ and optimized for nephron lineages ^37^. Biological replicates were generated from different iPSC batches. At 70-80% confluency iPSCs were rinsed with 1xPBS (Thermo Fisher Scientific, 10010049), isolated using TrypLE Select Enzyme (Thermo Fisher Scientific, 12563011) for 6 minutes at 37°C, enzymatic reaction neutralized in Essential 8 media, and cells collected and resuspended in Essential 8 media with 10 µM Y-27632. 10,000 cells were plated per 12-well plate well onto 5% biolaminin 521 LN and differentiation started 6 hrs post plating in TeSR-E6 (Stem Cell Technologies, 05946) with CHIR99021 (Tocris, 4423). Thereafter, media was supplemented with CHIR99021 for 5 days. CHIR99021 concentrations were optimized for each cell line. At day 5, media was changed for TeSR-E6 with 200 ng/mL FGF-9 (R&D Systems, 273-F9) and 1 µg/mL Heparin (Millipore Sigma, H4784). At day 7, cells were isolated using TrypLE and resuspended in TeSR-E6 with 10 µM Y-27632. 200,000 cells were seeded into round-bottom non-adhesive 96-well plate wells and allowed to self-aggregate. 3D organoid aggregates were manually transferred to transwell filters (Corning, 3450; Stem Cell Technologies, 100-1026), pulsed with TeSR-E6 with CHIR99021 for 1 hour and thereafter in TeSR-E6 with 200 ng/mL FGF-9 and 1 µg/mL Heparin until day 12. From day 12 onward, organoids were cultured in TeSR-E6 alone.

### DNA constructs

For *ROSA26*/*ROGI1* locus targeting, gRNA-encoding oligos were cloned into *pSpCas9-GFP* as described previously ^60^. Plasmids were screened via Sanger sequencing using the LKO.1 5’ primer. For insertion of *TagBFP-2A-Wnt3a/WNT9B* into *ROSA26*, *TagBFP* was PCR-generated from an in-house construct (*pTH11*); 2A was added downstream through two PCR rounds using sequential reverse primers. *Wnt3a* cDNA was PCR-generated from +Wnt3a-SO gDNA ^32^. *WNT9B* isoform-201 cDNA was acquired as a synthetic gene fragment following human codon GC-content optimization (Twist Biosciences). *TagBFP*/*WNT* fragments were assembled into KpnI-linearized *pHDR-ROSA26-MCS* vector established previously (FG’s thesis, gift from Jamie Davies). For insertion of *mCherry^Surface^-2A-Cdh1/Cdh3* into *ROGI1*, the *EF1α* promoter was PCR-generated from *pHDR-ROSA26-MCS*, mCherry^Surface^-2A from an in-house construct (*pMMP250-Cdx2*), and *Cdh1/Cdh3-pA* from E-cell/P-cell gDNA ^61^. *EF1α*/mCherry^Surface^/*Cdh* fragments were assembled into BamHI-linearized *pHDR-ROGI1-MCS* vector established in-house based on published reports ^44^. All PCRs were performed using Q5 polymerase master mix (NEB: M0492S); primer sequences are in the *key resources table*. Assemblies were performed using 30fmol vector, 50fmol per insert, HiFi DNA assembly reaction mix (NEB: E2621L), in 20μl final volumes at 50°C for an hour. Volumes corresponding to 2fmol of vector were used to transform Stellar competent bacteria via 42°C heat-shock for 30-35 seconds; 9 parts SOC medium were added to 1 part bacteria, transformants were outgrown at 37°C, 250rpm for 40 minutes, and one-tenth of the outgrowth was plated on Luria-Bertani agar plates containing 100μg/ml ampicillin. Four-to-six colonies were expanded in 3-4ml ampicillin-supplemented broth and purified plasmids were screened via restriction digestions and full plasmid sequencing (Primordium labs).

### Cell engineering

One-microgram *pSpCas9-gRNA* and 3μg *pHDR-ROSA26/ROGI1* constructs were added into 125μl OptiMEM together with 5μl P3000 reagent. The mixture was added into a pre-vortexed solution containing 125μl OptiMEM and 5μl Lipofectamine3000 reagent (ThermoFisher: L3000001). Solutions were incubated on site for 15 minutes. HEK-293 6-well cultures at 70-80% confluence were gently washed in PBS, replenished with 1ml OptiMEM, and transfection solutions were decanted drop-wise while rocking the plate. Transfectants were incubated at 37°C, 5% CO_2_ for 6 hours, and replenished with culture medium. Two days later, cultures were refed with medium containing 10μg/ml puromycin (*pHDR-ROSA26-WNT*) or 800μg/ml G418 (*pHDR-ROGI1-Cdh*). Drug-supplemented medium was replenished every 2-3 days for 12-14 days total, whereupon 1μM doxycycline was added for 24 hours. Cells were sorted on the basis of mCherry+TagBFP+ profile, cultured for 5 days, and re-sorted on the basis of mCherry+TagBFP-profile.

### Evaluation of organizer doxycycline tuneability

One-hundred-thousand cells were seeded in 12-wells. The next day, cultures were refed with fresh medium supplemented with 10-fold serial dilutions of doxycycline. Twenty-two hours after supplementation, cultures were harvested, resuspended in 0.6ml PBS-2% FBS, filtered through flow cytometry tube cap filters and analyzed on a BD FACSAria II cytometer.

### Synthetic organizer production

On organoid differentiation day 11, HEK cultures were harvested and cell concentrations determined using a Countess II FL automated cell counter (Invitrogen). Five-thousand cells per spheroid times desired number of spheroids were pelleted and resuspended in doxycycline-supplemented medium (50μl per 5×10^3^ cells). Fifty microliters were decanted in Nunclon Sphera 96U-well round bottom plate wells (Thermo Scientific 174929), the plate was centrifuged at 100g for 2 minutes, and placed in a 37°C, 5% CO_2_ incubator overnight.

### mRNA-sequencing and data analyses

Samples were prepared and purified according to the RNeasy Mini Kit (Qiagen, 74104). For iPSCs, one well from a 6-well plate was collected prior to kidney organoid differentiation. Whole organoid samples were collected for differentiation days 10 and 12 (2 organoids per biological replicate). Organoid regions adjacent to synthetic organizers were surgically isolated using a sichel knife (Fine Science Tools, 10073-14), and subsequently dissociated and purified sorted for DRAQ5+, DAPI-, and mCherry-. Wnt3a-SOs and control-SOs were cultured as 5,000 cell aggregates in a 96-well U-bottom plate for 24 hours before collection. Purified RNA was sent to Novogene for paired end sequencing. Reads were aligned to the hg38 Ensembl 105 annotation using star version 2.7.10a ^62^. A noise-reduction filter was applied to keep genes in which the maximum count in or at least one sample was greater than or equal to 10. Partek Flow version 10.0 was then used to normalize raw feature counts for Wnt3a-SOs and control-SOs following the transcripts per million (TPM) normalization method. Separately, iPSC, organoid, and previously published human kidney datasets ^11^ were normalized using the DESeq2 median ratio method ^63^. One-sided Welch’s tests were used to perform statistical analyses using *t.test()* with the parameter *var = F* in R. Data visualization was made using *ggplot2()* in R.

### Single-cell RNA sequencing and analyses

Microdissected organizer-coupled organoid regions were dissociated with Accumax (STEMCELL Technologies 07921) for 20-30 minutes, neutralized with double the amount of autoMACS buffer, filtered through flow cytometry cap filters and sorted for mCherry-, DAPI-DRAQ5+ live cells. Cells were processed for the split-seq Evercode WT v2 single-cell RNA-sequencing platform (Parse Biosciences, ECW02030) as described previously ^37^. Samples were prepared for paired-end 150nt sequencing using NovaSeq X Plus. Fastq files were demultiplexed, quality controlled, and aligned (hg38 Ensembl 105 annotation), using the *splitpipe* workflow (Parse Biosciences). High-quality cells (>500 genes, <35% mitochondrial gene content) were kept using the Seurat 5.0 package. Samples were integrated using the Seurat integration function. Differentially expressed genes (>25% cells, >0.25-fold change) were identified using the *FindAllMarkers* function. Clusters were identified based on published nephron and interstitial markers ^15,16,18^. Day 10, 12, and 14 kidney single cell RNA sequencing was performed as described previously ^37^ Developing kidney single cell profiles (week 14 and 17 post conception) were obtained from published datasets ^16,18^ and integrated to generate the *in vivo* nephrogenesis framework ^37^.

### Spatial transcriptomic analyses of developing kidneys

Whole kidneys were fixed at 4°C for 4 hours in 4%PFA (Electron Microscopy Sciences, 15710), thoroughly washed with PBS, and embedded in OCT. 10um sections were prepared for Xenium and GenePS platforms as per manufacturers’ recommendations (10X Genomics, Spatial Genomics). Gene probesets (249-250 genes) were designed to capture all kidney populations at week 16 and the nephrogenic gene expression program. Nephrogenic niches and stages were identified based on morphologies and gene expression patterns. Transcript positions were quantified using R-studio and Fiji to examine transcript coordinates relative to the collecting duct tip and stalk using *WNT11* and *WNT9B* as markers.

### RNA scope and quantification of WNT4 polarity

Whole kidneys fixed and prepared as for spatial transcriptional analyses were processed for RNA scope as previously described ^18^. RNAscope images of 8 renal vesicle-staged nephrons stained for *WNT4*, *WNT11*, *WNT9B*, *KRT8* and DAPI were processed on ImageJ/FiJi. The nephron’s lumen center and *WNT11*/*WNT9B* boundary were marked with the *point* tool. *WNT11, WNT9B* and *WNT4* puncta were registered using the *multi-point* tool. Lines were drawn between the nephron’s lumen center point and the *WNT11*/*WNT9B* boundary using the *straight-line* tool, and between the nephron’s lumen center point and *WNT* puncta using a custom macro. The angle of each line relative to the lumen center was recorded, summarized and averaged for all nephrons, binned and graphed in a radar plot using a custom Python script.

### Quantification of spatial transcriptional data in the NPC population and PTA along an adjacent-distant axis

Spatial transcriptomics data from capturing a total of 9 CD tips and nephron progenitors with associated PTAs, from week 13 and week 16 kidneys, were analyzed for expression of *PCDH15*, *MEOX1*, *ELAVL4*, and *FGF8*. The distribution of transcripts was graphed in relation to the surface of each *WNT11*^+^ CD tip. Pretubular aggregates were defined based on morphologies (DAPI) and the collective view offered by expression of PCDH15, MEOX1, ELAVL4, FGF8, WNT4, LHX1, JAG1, EMX2, GATA3, POU3F3, HNF1B, LEF1, PDGFRB, in relation to the WNT11/WNT9B transition in the CD. Transcript distribution was plotted in PTAs for WNT4, FGF8, LHX1, EMX2, JAG1, MEOX1, and PCDH15 with respect to the forming lumen in the PTA and the WNT11/WNT9B transition point. The angle of each line relative to the lumen center was recorded, summarized and averaged for all nephrons, binned and graphed in a radar plot using a custom Python script.

### Quantification of Morphogenesis

Organizer-coupled HNF4A::YFP organoids were imaged via widefield fluorescence microscopy on Day18 and processed on ImageJ/FiJi. The center of the mCherry+ organizer was marked with the *point* tool, arrows were drawn from YFP+ tubules to YFP-tubules in clearly polarized nephrons showing elongated morphologies around the organizer, and axial paths were extrapolated using the *straight-line* tool. If paths passed through the organizer, they stopped at the organizer’s end; if not, they stopped when they could not converge with any other extrapolated path. Intersections were registered using the *multi-point* tool. XY coordinates of the organizer’s center and intersections were extracted and the Euclidean distance between each intersection and the organizer’s center calculated and averaged per organoid.

### Immunostaining & Clearing

Kidney organoids and mouse kidney explants were fixed for 20 min on ice in 4% PFA. Samples were permeabilized and blocked using PBS with 1.5% SEA Block (Thermo Fisher Scientific, 37527X3) and 0.1% TritonX100 (EMD Millipore, 1.08643) at 4°C with gentle movement for 1 hour. Samples were stained with primary antibodies in blocking solution overnight. Stained samples were washed for at least 3 hours through several rounds of PBS-0.1% TritonX100 washes, and stained with secondary antibodies in blocking solution overnight. PBS washing steps were repeated the next day and nuclei were counterstained using PBS-0.1% TritonX100 containing 1 µg/mL Hoechst 33342 (Thermo Fisher Scientific, H3570) for 25 minutes. For widefield microscopy, samples were washed for 1 hour in PBS and mounted on slides with a glass coverslip in Immu-Mount (Thermo Fisher Scientific, 9990402). For confocal microscopy, organoids were cleared by gradual dehydration in and placed in BABB ^16^.

### Confocal Imaging & Volume Analysis

Optically-cleared organoids were mounted in 20mm-diameter FastWell frames (Grace Bio-Labs, #664113) loaded with 190μl benzyl alcohol-benzyl benzoate (BABB). A coverslip shard was put on top of the transwell to ground the organoid, another round coverslip was used to seal the FastWell, and the sealed chamber was nail polish-glued to a Superfrost plus microscope slide. Organoids were imaged on a Leica SP8X confocal laser scanning microscope by stitching 20 Z-stack tiles captured using 40X oil lens (512×512 resolution). For thicker organoids a 25X water long-working distance lens was used. Imaging files extracted as TIFs were converted to IMS format using Imaris file converter and processed on Imaris imaging analysis software. Organizer surfaces were rendered from manually drawn regions on the basis of strong CDH3 or faint TFAP2A signal; a positive mask was extracted from the surface (organizer channel), and negative masks subtracting the organizer from all other channels were produced. Surfaces from resulting organizer-subtracted channels were rendered with a minimum 2000μm^3^ volume threshold, and manually curated by deleting objects stemming from the mesenchyme, organizer leftover signal, and the autofluorescent transwell. Statistics and surface galleries were extracted via the *Vantage* tab. 3D renders were visualized using the Blend option and images captured via the *Snapshot* tab.

### Statistics

The exact sample size (n) and statistic test employed for each dataset are detailed in the *key numbers table*. Repeated measurements were performed when different nephrons from the same organoid were quantified, when different immunostain intensities from the same nephron region were extracted, and when different metrics from the same image were extracted. Unless otherwise noted, data were assumed to be normally distributed. Significance and effect sizes are noted on figure panels (*P*: P-value, *f.c.*: fold-change, *D*: Cohen’s d calculated at https://www.socscistatistics.com/effectsize/default3.aspx). Means and standard deviations are described in text. One-tailed t-tests were performed if the variable tested was expected to exert a directional change.

---

**Figure S1.**
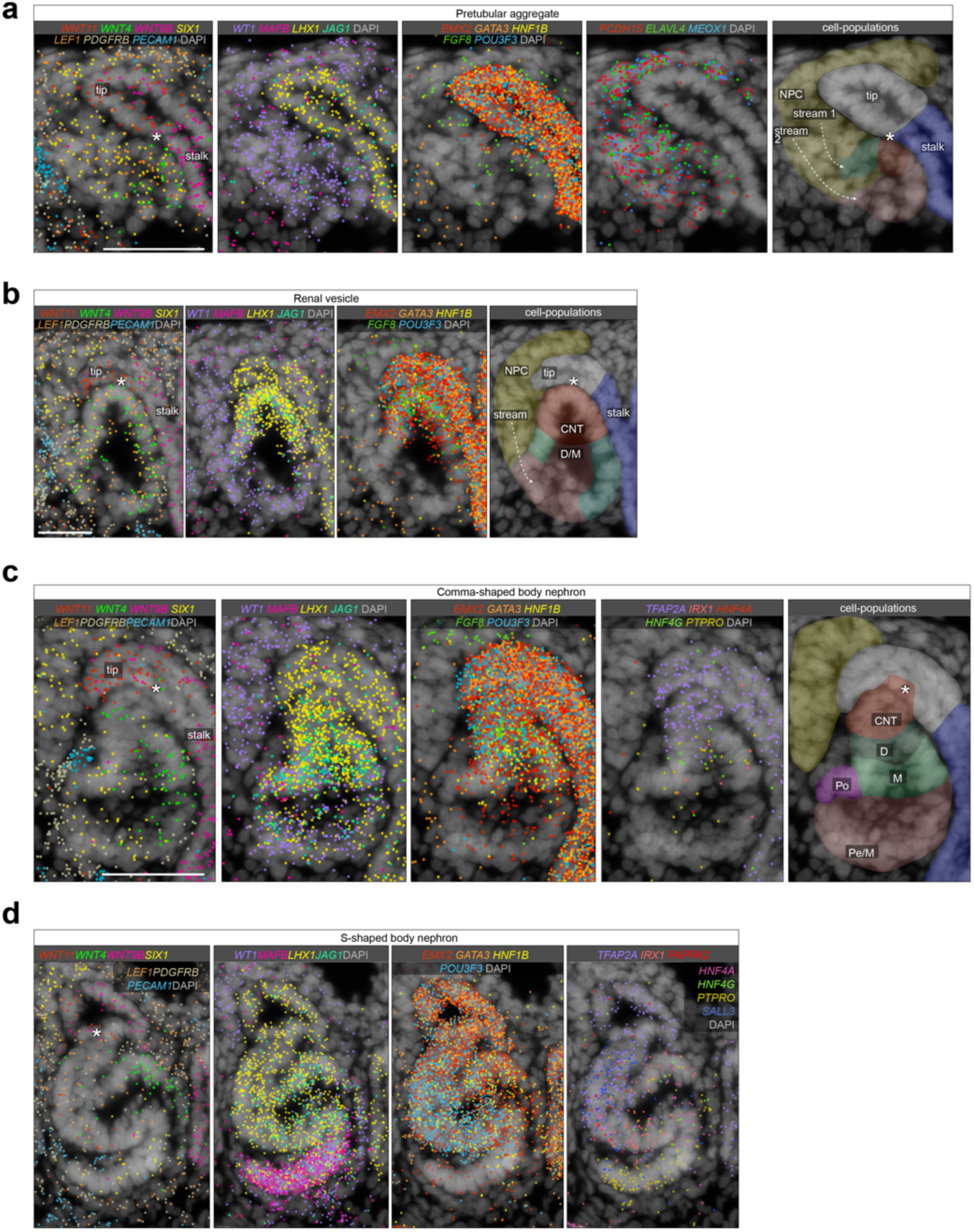
Spatial transcriptomic data of human developing nephrons (in support to Figure 1a-h). **a,** Xenium spatial transcriptomic data examples of developing human nephrons at the pretubular aggregate, renal vesicle, comma-shaped body, and S-shaped body nephron stages with genes shown as in Figure 1.

**Figure S2.**
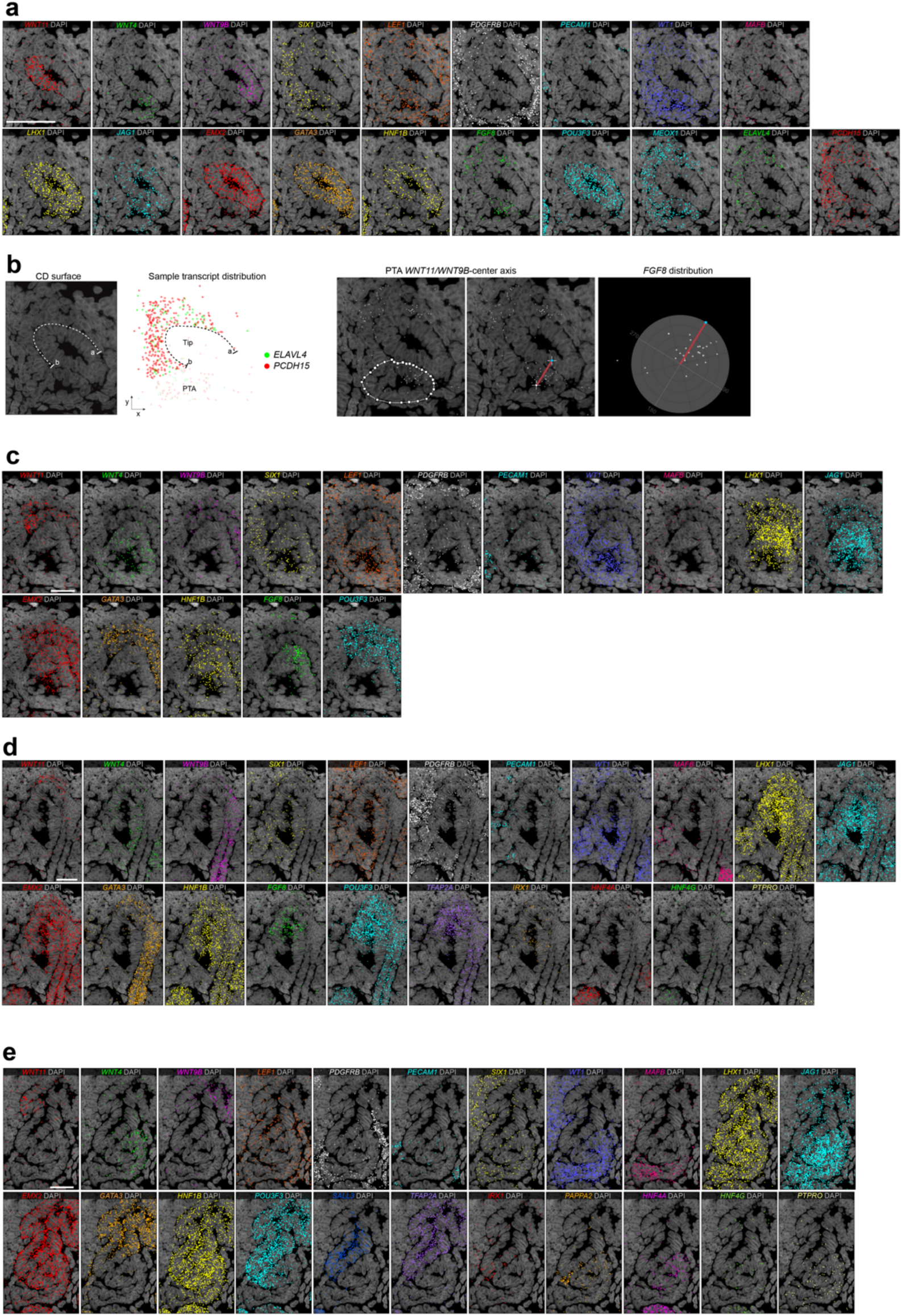
Spatial transcriptomic data breakdown and analysis (in support to Figure 1a-h). **a,** Single-channel breakdown of overlayed spatial transcriptomic data at the pretubular aggregate stage shown in Figure 1c. **b,** Method for quantifying distances and distribution between various gene transcripts (dots) and the CD surface (dotted white line) in nephron progenitors and pretubular staged nephrons from 13.5 and 16.5 week human developing kidney sections, related to Figure 1d, **c-e,** Single channels for renal vesicle, comma-shaped, and S-shaped body stages of nephrogenesis.

**Figure S3.**
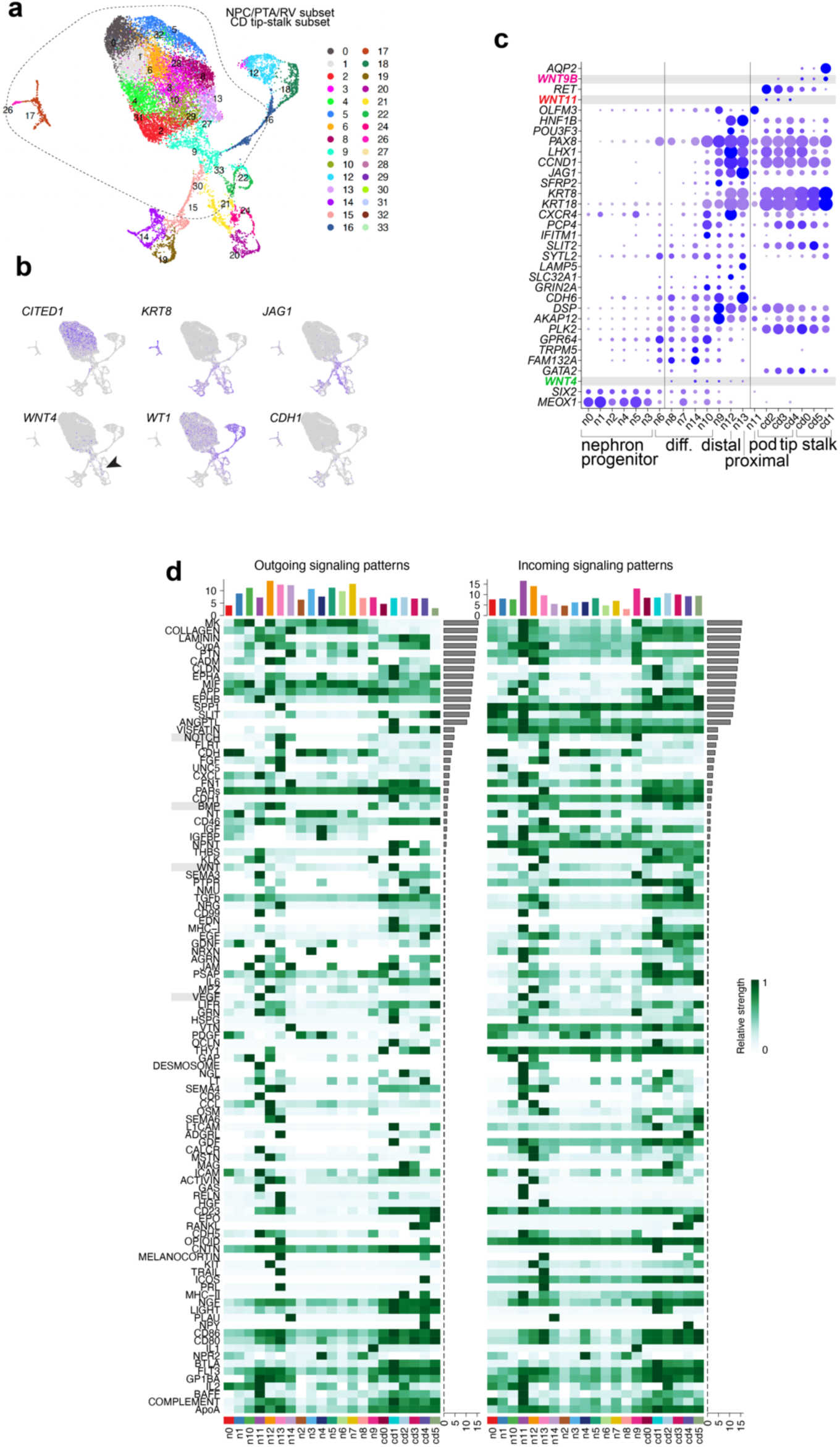
Human developing kidney single-cell RNA sequencing analyses (in support to Figure 1i-k). **A.** Single-cell RNA sequencing UMAP showing clustering of nephrogenic and collecting duct. **b,** Expression of key nephron markers in the UMAP shown in (a), showing self-renewing progenitors in *CITED1*, epithelializing tubular precursors in *KRT8*, developing renal vesicle and medial precursors in *JAG1*, differentiating progenitors in *WNT4*, podocytes and nephron progenitors in *WT1*, and epithelial tubule populations in *CDH1*. **C.** Expression of the collecting duct stalk and tip markers, and genes enriched in self-renewing nephron progenitors, differentiating nephron progenitors, distal precursors, proximal precursors, and podocyte precursors, used to annotate clusters shown in Figure 1i-k. **d.** Ligand-receptor pairing analyses using CellChat to predict cell signaling in the early collecting duct and nephron lineage.

**Figure S4.**
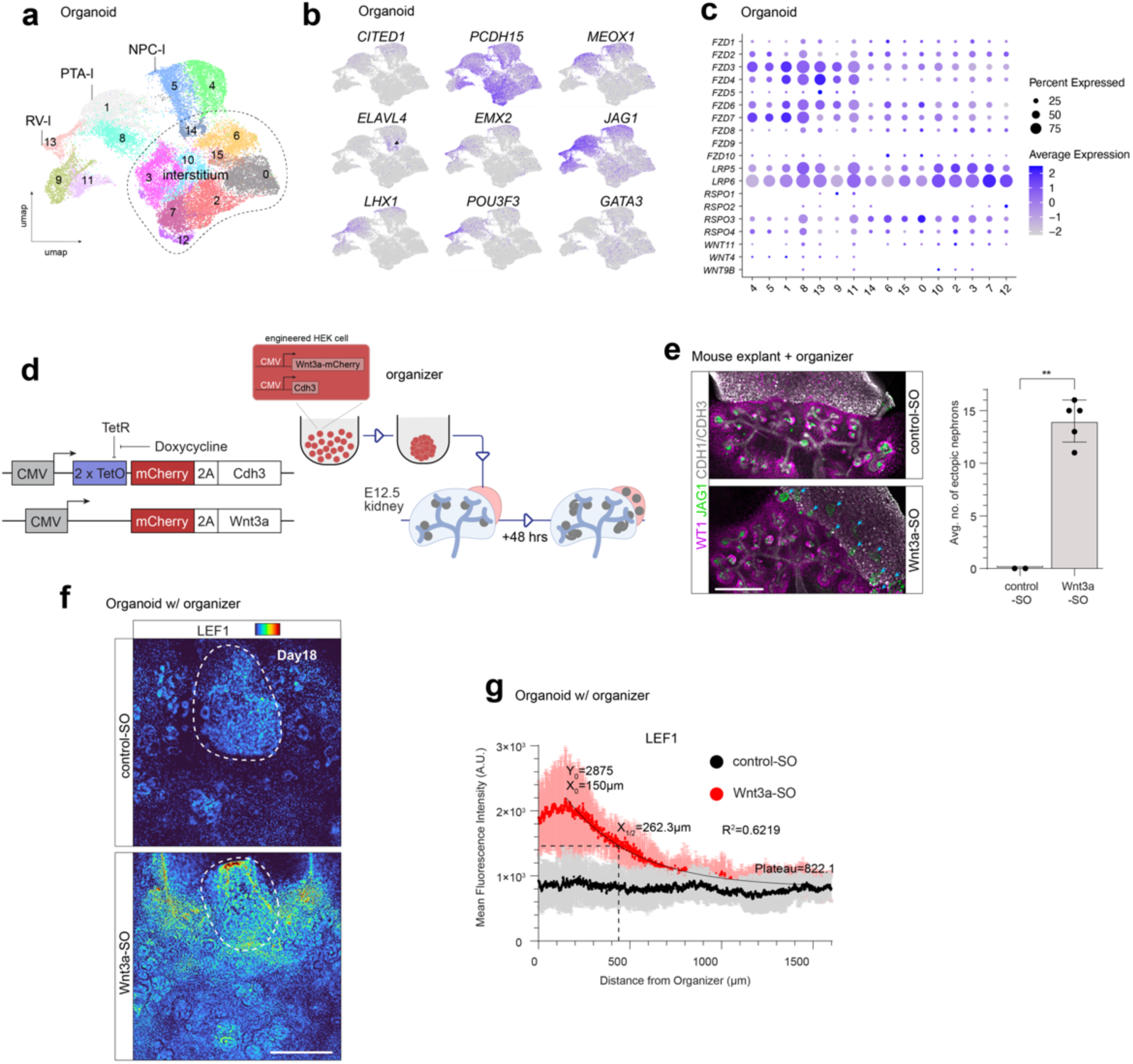
Absence of Wnt/β-catenin interactions in human kidney organoids and gradient reconstitution via synthetic organizers (in support to Figure 2d-j). **a,** UMAP plots of Day10-14 human kidney organoids identify nephrogenic cells transitioning from progenitor-like to pretubular aggregate-like to renal vesicle-like profiles in the presence of interstitium. **b,** Gene expression of key markers enriched in self-renewing progenitors (*CITED1*), early differentiating progenitors (*PCDH15, MEOX1, ELAVL4*), the committed renal vesicle (*JAG1, LHX1*) and distal precursors (*EMX2, POU3F3, GATA3*) in the UMAP shown in (a). **c,** Expression of WNT ligands implicated in kidney development, Frizzled1-10 receptors, LRP5/6 co-receptors, and RSPO1-4 potentiators of Wnt/β-catenin activity across nephrogenic and interstitial clusters of kidney organoids. **d,** Outline of supplemental experiment testing the ability of synthetic organizers to influence nephron development programmes, determined by inducing ectopic nephrogenesis in mouse kidney explants. **e,** Immunostain examples and quantifications of ectopic JAG1+ nephrons forming in the mouse metanephric mesenchyme adjacent to WNT organizers, but not in contact with the collecting duct. **f-g,** Immunostained Day18 organoids coupled with constitutive Wnt3a organizers or controls revealing a long-range β-catenin activity gradient as determined by LEF1 intensity quantification over increasing distance from the organizer.

**Figure S5.**
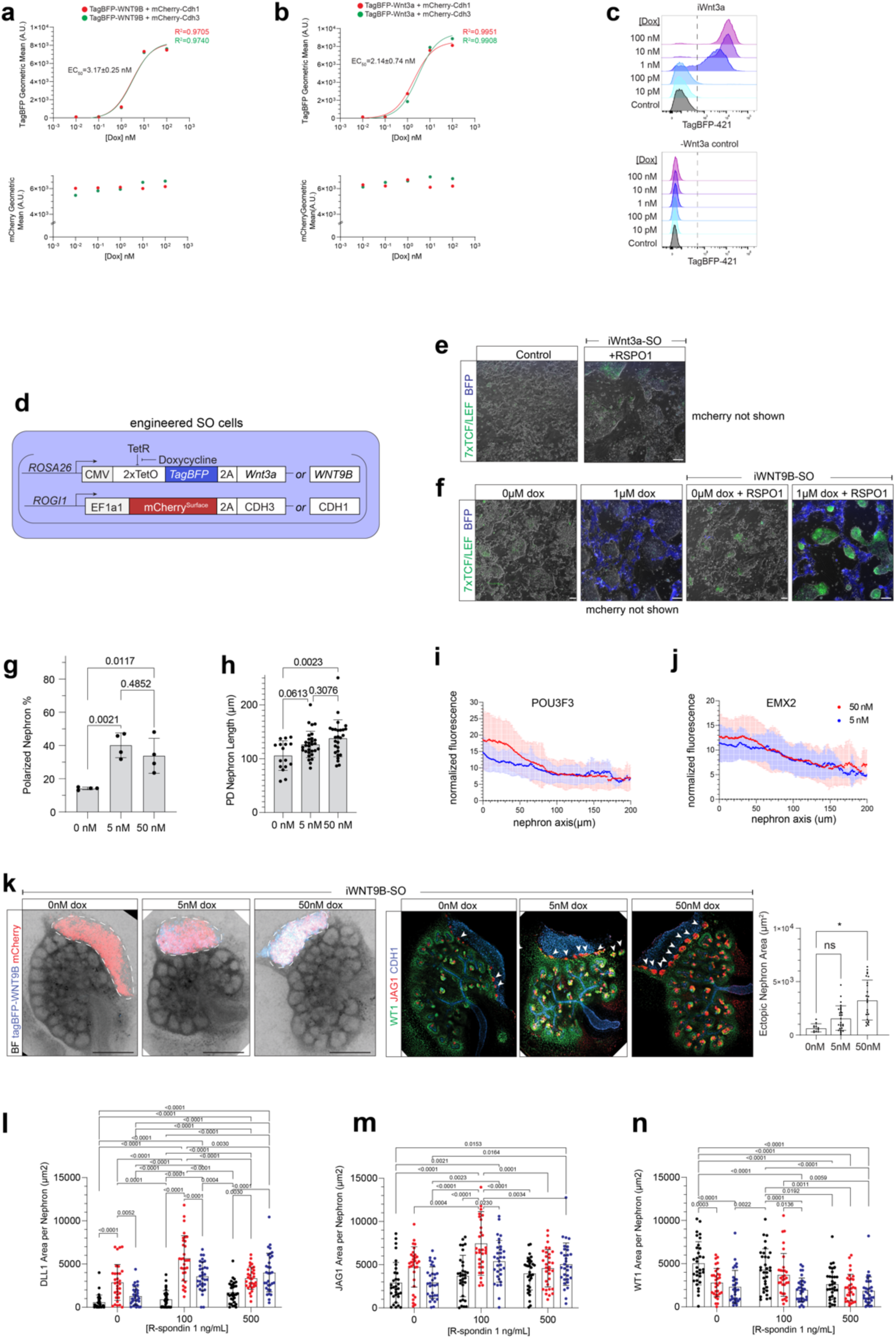
Nephron polarization driven by bioengineered dox-tunable WNT secreting synthetic organizers (in support to Figure 3). **a-b,.** Dose-response curves showing geometric mean fluorescence intensity of the WNT-reporting TagBFP module (top panel) and the cadherin-reporting mCherry^Surface^ module (bottom panel) against 10-fold incremental increases in doxycycline concentration. Dose-response curves show results for iWNT9B organizer and iWnt3a organizer versions harboring different cadherin modules. **c,** TagBFP fluorescence histograms of iWnt3a organizer cells after 24h exposure to indicated doxycycline concentrations. **d,** Synthetic organizer blueprint driving mCherry^Surface^-reported *Cdh3* expression from the *ROGI1* locus constitutively, and *TagBFP*-reported WNT expression from the *ROSA26* locus under doxycycline induction. **e-f,** Live fluorescence imaging of co-cultures between iWnt3a or iWNT9B cells with β-catenin reporter eGFP mouse embryonic stem cells, revealing differential signaling response and RSPO1 dependency depending on WNT ligand specificity, respectively. **g,** Percentage of nephrons showing polarized EMX2/POU3F3 phenotype in Day 14 organoids, for renal vesicles around and 2-3 layers away from iWnt3a-SOs, as shown in Figure 3c. **h.** Length of polarized nephrons shown in (g). **i-j,** Quantification of EMX2 and POU3F3 intensities along the organizer adjacent-distant axis in iWnt3a-SO polarized nephrons, as shown in Figure 3e. **k,** Mouse kidney explant experiment showing ectopic nephrogenesis induction by iWNT9B-SOs, and a dox/WNT tunable increase in ectopic nephron area. **l-m,** Graphs showing immunofluorescent intensity quantification of DLL1, JAG1, WT1 as relating to Fig.3k-m with p-values.

**Figure S6.**
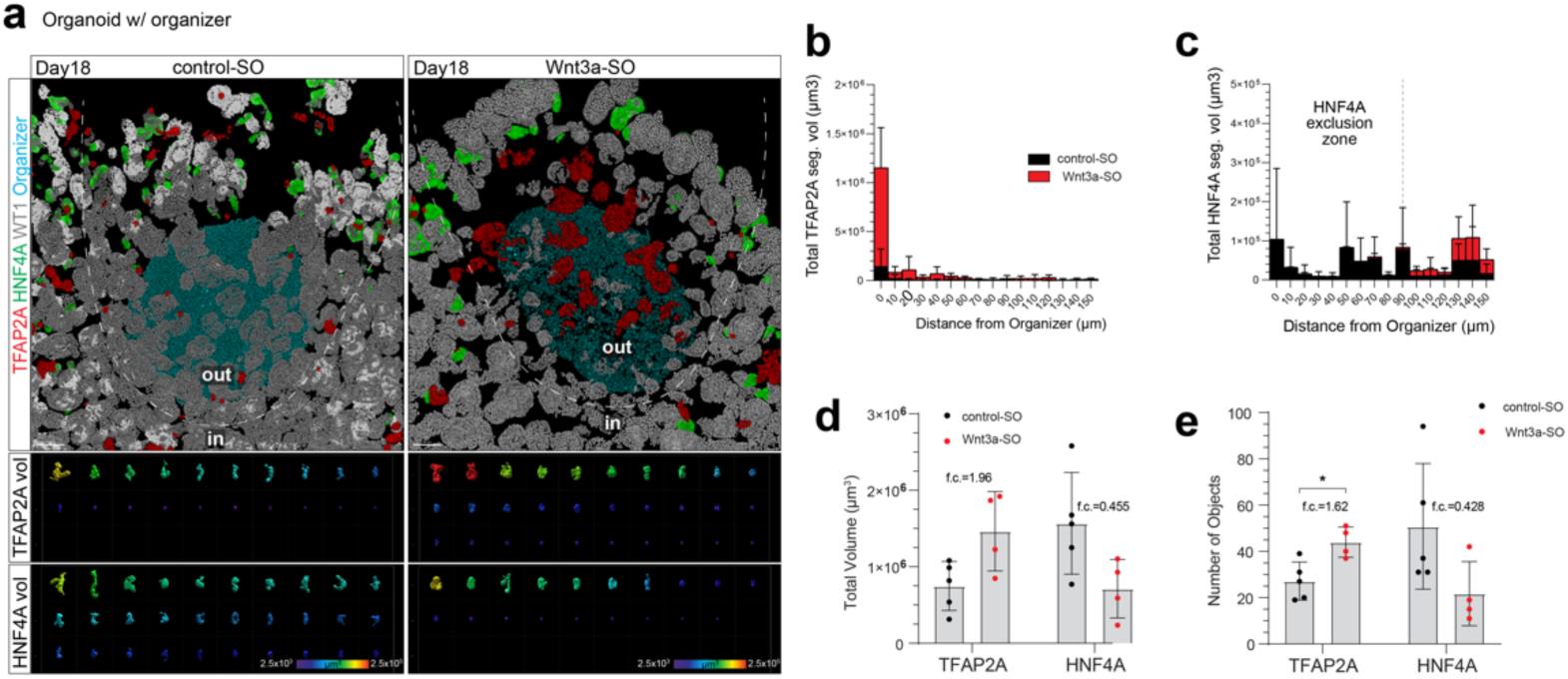
Distal nephron segments expand at the expense of proximal segments in Day 18 organoids coupled to constitutive WNT synthetic organizers (in support to Figure 4a-e). **a,** 3D render snapshots of Day18 organoids coupled with control spheroids or constitutive *Wnt3a* organizers on Day12, immunostained against TFAP2A distal tubules, HNF4A proximal tubules and WT1 podocytes. Bottom panels show top 30 TFAP2A and HNF4A objects descending by volume, for nephrons located underneath the organizer and towards the organoid’s periphery (under-and-out, above dotted line). **b-c,.** Distribution of TFAP2A (b) and HNF4A (c) volumes relative to the distance from the constitutive *Wnt3a* organizer in 10μm increment bins for nephrons in the under-and-out position. **d-e,** Total volumes (d) and number counts (e) of TFAP2A and HNF4A segments in Day18 organoids coupled with control or constitutive *Wnt3a* organizers, in under-and-out nephrons.

**Figure S7.**
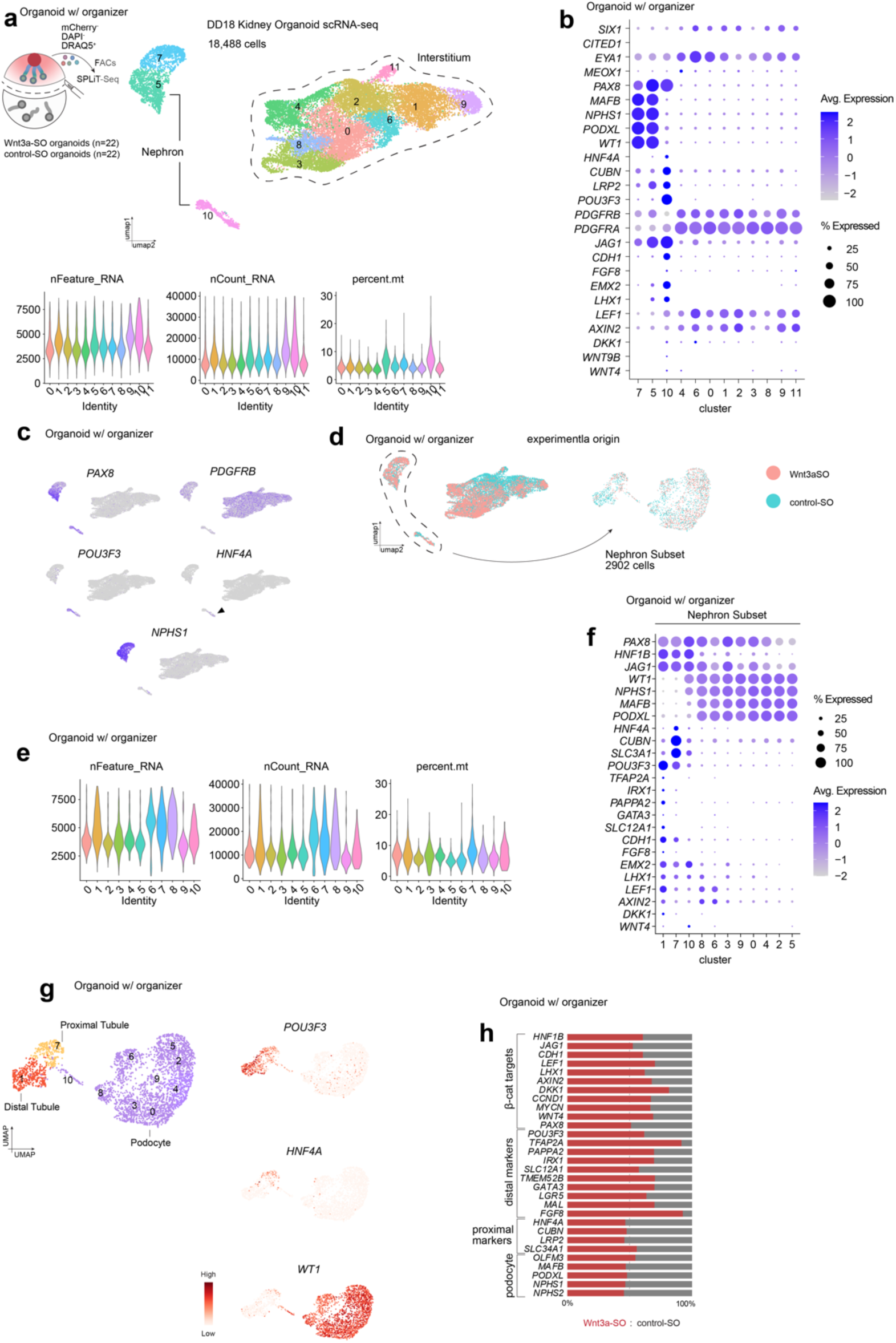
Activation of distal tubule development in Day18 organoids coupled with WNT synthetic organizers (in support to Figure 4). **a,** Overview of single-cell RNA sequencing for nephrons near signaling centers following exclusion of mCherry+ HEK-293 cells and selection of DAPI-DRAQ5+ live cells (n=22 organoids). UMAP shows segregated interstitium and nephron clusters; violin plots show sequencing quality control metrics (gene reads per cell, total reads per cell, mitochondrial content per cell) across annotated clusters **b,** Expression of nephron progenitor (SIX1, CITED1), podocyte (MAFB, NPHS1, PODXL, WT1), proximal (HNF4A), distal (POU3F3, EMX2) and medial (JAG1, FGF8) precursor markers, interstitium-enriched genes (PDGFRA, PDGFRB), as well as kidney WNT ligands (WNT9B, WNT4) and downstream effectors (DKK1, AXIN2, LEF1) across annotated clusters. **c,** Expression of kidney (PAX8), interstitium (PDGFRB), podocyte (NPHS1), proximal precursor (HNF4A) and distal precursor (POU3F3) markers in the UMAP. **d.** Overlay plot of origin UMAP showing whether data points originate from control or +cWnt3a organizer-conjugated samples, and further analysis within the nephron clusters. **e,** Quality control metrics within subet nephron-only clusters. **f,** Expression of β-catenin targets, podocyte, proximal, medial, and distal lineage marker expression across nephron-only clusters. **g,** UMAP plots showing podocyte, proximal tubule, and distal tubule populations based on segregated expression of *WT1*, *HNF4A*, and *POU3F3* respectively within nephron-only UMAPs. **h,** Percentage of transcript reads coming from WNT-vs-control-conjugated organoids for indicated genes, for the total sum of reads for each gene.

**Figure S8.**
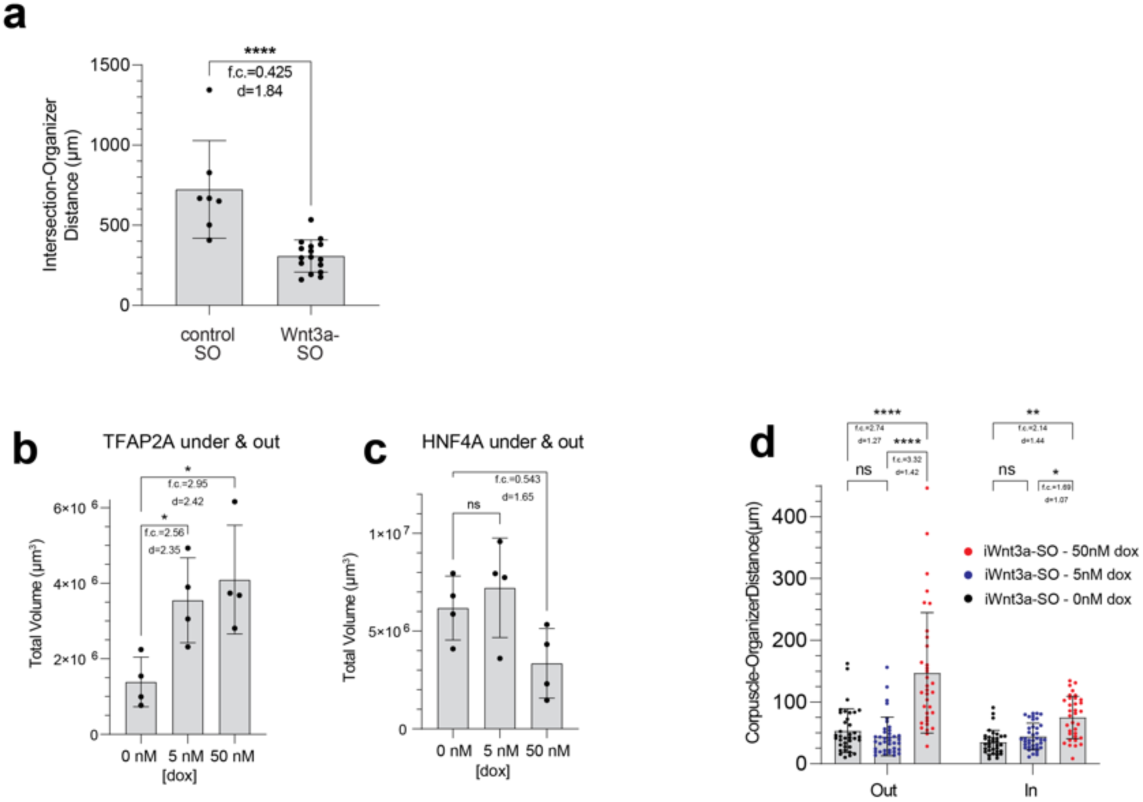
Morphoalignment and axial patterning in organoid nephrons enabled by WNT secreting organizers (in support to Figure 4). **a,** Distance between tubule path intersections and the organizer’s center per organoid. **b-c,** Total TFAP2A (d) and HNF4A (e) volumes for nephrons in the under-and-out position in iWnt3a-coupled Day18 organoids as in Figure 4i. **d,** Distances between organizer peripheries and nearest renal corpuscles located towards the organoid’s periphery (out) or center (in), for iWnt3a-coupled Day18 organoids at different dox dosages.

## Supplementary Tables

**Supplementary Table 1.** Differentially expressed genes within single-cell RNA sequencing dataset clusters of developing human kidneys at 14-17-week post conception.

**Supplementary Table 2.** Bulk RNA sequencing normalized gene expression comparisons between 9.5-21-week post conception developing human kidneys and Day 10-12 kidney organoids.

**Supplementary Table 3.** Bulk RNA sequencing normalized gene expression in control and +Wnt3a-SOs and isolated HEK cells, sheets A and B, respectively.

**Supplementary Table 4.** Differentially expressed genes within single cell RNA sequencing dataset clusters of Day 10-14 kidney organoids.

**Supplementary Table 5.** Bulk RNA sequencing normalized gene expression comparisons between standalone Day 14 kidney organoids and kidney organoids coupled with control or constitutive *Wnt3a* organizers.

**Supplementary Table 6.** Differentially expressed genes between single cell RNA sequencing datasets of Day 18 kidney organoids coupled with control or constitutive *Wnt3a* organizers, for nephron-like clusters.

**Supplementary Table 7.** Differentially expressed genes between single cell RNA sequencing datasets of Day 18 kidney organoids coupled with control or constitutive *Wnt3a* organizers, for all clusters.

